# Microglial-Mediated Prevention of Axonal Degeneration in the Injured Spinal Cord: Insights from an *In Vivo* Imaging Study

**DOI:** 10.1101/2024.04.13.589343

**Authors:** Wanjie Wu, Yingzhu He, Yujun Chen, Yiming Fu, Sicong He, Kai Liu, Jianan Y. Qu

## Abstract

Microglia, the primary immune cells in the central nervous system, play a critical role in regulating neuronal function and fate through their interaction with neurons. Despite extensive research, the specific functions and mechanisms of microglia-neuron interactions remain incompletely understood. In this study, we demonstrate that microglia establish direct contact with myelinated axons at Nodes of Ranvier in the spinal cord of mice. Under normal physiological conditions, microglia-node contact occurs in a random scanning pattern and is associated with neuronal activity. However, in response to axonal injury, microglia rapidly transform their contact into a robust wrapping form, preventing acute axonal degeneration from extending beyond the nodes. This neuroprotective wrapping behavior of microglia is dependent on the function of their P2Y12 receptors, which may be activated by ATP released through axonal volume-activated anion channels at the nodes. Additionally, voltage-gated sodium channels (NaV) contribute to the interaction between nodes and glial cells following injury, and inhibition of NaV delays axonal degeneration. Through *in vivo* imaging, our findings reveal a neuroprotective role of microglia during the acute phase of spinal cord injury, achieved through a novel form of neuron-glia interaction.

## INTRODUCTION

Spinal cord injury (SCI) is a devastating condition with profound, life-altering consequences, yet currently lacking a cure. The study of axon damage and regeneration in SCI has been impeded by the absence of appropriate *in vivo* imaging tools capable of visualizing undisturbed cellular processes within the spinal cord. In this work, we utilize multimodal microscopy and optical clearing technology for minimally invasive *in vivo* imaging study of the pivotal role played by microglia in axonal degeneration following SCI. Microglia is the main resident immune cells in the central nervous system (CNS) and known to play a critical role in brain development, homeostasis, and neurological disorders^1–3^. In the healthy CNS, microglia exhibit a ramified morphology, and their motile processes continuously survey the surrounding microenvironment^4,5^. Upon disruption of CNS integrity, microglia is capable to rapidly respond by adapting themselves to a morphologically and biochemically distinct activated state^6–9^. Recent research has provided increasing evidence for microglial direct contact with neurons, which modulates neurogenesis, synaptic plasticity, neuronal firing, as well as neurodegeneration and regeneration in response to external insults^10,11^.

Direct physical contacts between microglia and neurons establish a more efficient, precisely-controlled, and versatile form of regulating neuronal activities and fate, without the involvement of intermediate cells and soluble factors^10^. However, the direct membrane contact between microglia and neurons can be diverse and complex due to the large and polarized structure of neurons as well as the high-degree functional independence of their dendritic and axonal compartments^12,13^. In addition to the synaptic elements, which have long been the focus of microglia-neuron interaction^14,15^, specialized somatic microglia-neuron junctions have been identified in recent work and have been shown to protect neuronal functions^16^. Direct contact between microglia and axon has also been widely observed across species and is thought to support axonal function in the developing and adult CNS^10,11,17^. Specifically, the Nodes of Ranvier (NR), which are the short unmyelinated domains along myelinated nerve fibers, have recently been verified contacted by multiple glial cells and, especially, a preferential site for microglia-axon communication^18,19^. Microglia-axon contacts at NR have been shown durable, which contribute to remyelination following a demyelinating insult^18^. In our previous work, we observed that microglia recruit their processes to converge at NR, forming a wrapping contact in response to axonal injury in spinal cord^19^. Furthermore, NRs have been established as the initial disruption sites in a multiple sclerosis mouse model^20^, as well as the axonal sprouting sites during early axonal outgrowth after spinal cord injury^21^. The identification of microglial contacts at the nodal domain, as well as the involvement of NR as a special axon structure in CNS injury and repair, has led to the hypothesis that NR could potentially function as a pivotal hub, regulating axonal function and fate via its interaction with surrounding microglia.

Rodent models of spinal cord injury have been widely utilized to study axonal degeneration and regeneration^22,23^. In mouse models of spinal cord injury, it has been observed that axons undergo an acute bidirectional degeneration for a limited distance following the injury^21,24,25^. In the following days, the distal segments are completely destroyed via Wallerian degeneration while the proximal ends likely initiate a limited regeneration response. Despite significant efforts, the cellular and molecular mechanisms underlying the axonal response to injury remain incompletely understood^26–31^. The investigation of microglia-NR interactions may provide a new perspective for comprehensive understanding of axonal response to injury. Conventional methods for inducing spinal cord injury typically involve mechanical damage, such as transection, crush, or contusion^32^. Although these injury models mimic the clinical situation, they create a complex microenvironment around the lesion site that involves numerous variables working together to affect both anatomical and functional outcomes. Consequently, it is challenging to systematically examine the effect of a specific variable on the axonal response. A more precisely defined method of axonal injury, laser axotomy guided by two-photon imaging, allows to produce single axon injury at the target location^19,24,33–35^. *In vivo* two-photon imaging equipped with highly localized laser axotomy offers a flexible way to investigate microglia-node interactions in response to different well-controlled injury inputs^21,24,36–43^.

In a prior study, we developed an optically-cleared intervertebral window for in vivo spinal cord imaging without inducing microglial morphological activation ^19,44^. In this work, we employed this well-established intervertebral window to examine microglia-neuron interactions at NR. Through time-lapse two-photon excited fluorescence (TPEF) and stimulated Raman scattering (SRS) imaging, we observed that, under normal physiological condition, microglia contact nodes in a random scanning pattern, which is modulated by neuronal activity. Following axonal injury induced by laser axotomy, microglia rapidly establish a specialized wrapping contact by recruiting their processes to NR, effectively halting axonal acute degeneration before or at the wrapped nodes. Furthermore, we demonstrate that the protective wrapping behaviour of microglia is reliant on the function of its P2Y12 receptor. Additionally, inhibiting the axonal volume activated anion channels (VAACs) can disrupt the wrapping contact between microglia and NR, suggesting that purinergic signalling may regulate this contact. Finally, the involvement of sodium and potassium voltage-gated channels in the microglia-NR interaction following axon injury has been investigated in this work.

## Results

### Physiological microglia-axon interaction at NR in mouse spinal cord

To investigate microglia-axon interactions in real-time *in vivo*, we used a double crossed CX3CR1^+/GFP^/Thy1-YFPH transgenic mouse line to visualize microglia together with myelinated axons in spinal dorsal column. An optically-cleared intervertebral window was adopted to provide optical access to the spinal cord without introducing immune artifacts from surgical preparations^19^ (Fig. S1a). Through this window, we conducted *in vivo* TPEF and SRS imaging of the spinal cord, which revealed the structure of the NR, where the YFP axon was noticeable thinning and the myelin SRS signal was absent (Fig. 1a-c). By performing immunostaining of Caspr, which was located at the paranodes^18^, we confirmed that the SRS and YFP signal indeed identified the NR precisely (Fig. 1d). Three-hour time-lapse imaging with a 5-minute time interval showed microglial processes constantly surveying the surrounding microenvironment and dynamically contacting the NR (Fig. 1e, movie S1). Specifically, single microglial cell (microglia B) can contact multiple nodes (node 1 and 2) at the same time and single NR (node 1) allows multiple contacts from different microglial cells (microglia A and B). Most of the nodal contacts by microglia are intermittent and do not persist in a durable way (Fig. 1f-g). Among the observed 67 NR from 7 mice, 49 of them were contacted by microglia in a random scanning manner, with the duration of contact varying across different time points for each node. In addition, the remaining 18 nodes were observed to be continuously contacted by microglia within 3 hours. These *in vivo* findings present a contradiction to previous results based on an *in vitro* study, which suggested a preference for sustained microglial contact with nodes rather than a random scanning mode of contact^18^.

**Fig. 1.**
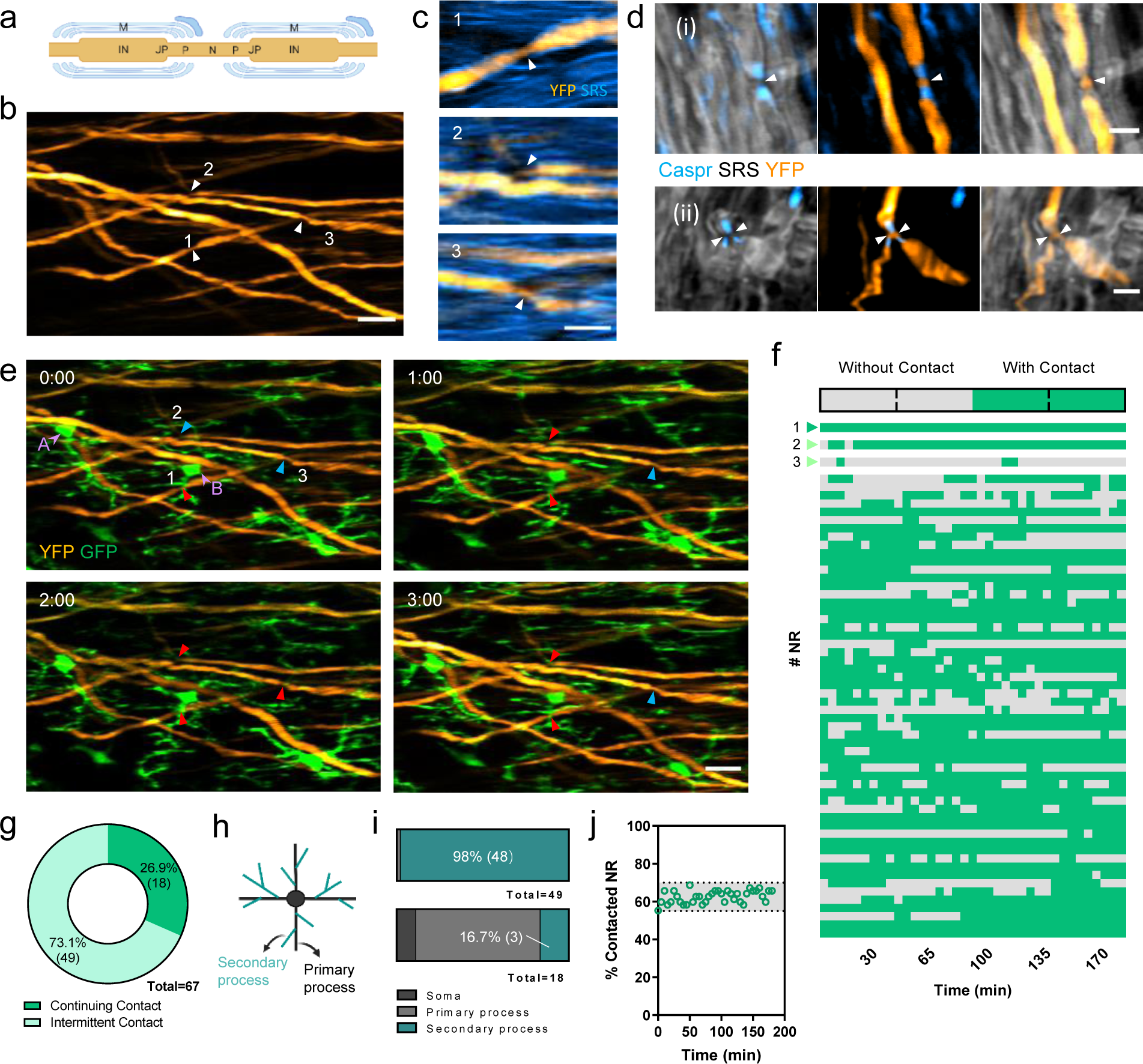
Microglial physical contact with Nodes of Ranvier (NR) in the normal spinal cord. (a) Morphologic components of a myelinated axon. M, myelin sheath; IN, internode; JP, juxtaparanode; P, paranode; N, node. (b, c) *In vivo* imaging of multiple NRs in mouse spinal cord. White arrowheads indicate the location of the same numbered NR in (b) and (c). Orange, YFP-labelled axons; blue, stimulated Raman scattering (SRS) image of myelin sheath taken at 2863.5 cm^-1^. (d) Immunofluorescent image of Caspr (blue), merged with SRS image of myelin sheath (gray) and YFP image of axons (orange) showing the position of normal node (i) and branching nodes (ii) (white arrowheads) in mouse spinal cord. (e) Three-hour time-lapse imaging of the nodes 1-3 shown in (b-c) together with surrounding microglia (green), showing dynamic microglia-node interaction. Red arrowheads indicate the occurrence of microglia-node contact while blue arrowheads indicate no microglial contact at the target nodes. Purple arrowheads show two microglial cells which contact the same node (Node 1) at the same time. Time is presented as hr: min. (f) Heatmap showing microglia-node contact events during three-hour time-lapse imaging. Each row represents a NR. Totally 67 nodes from 7 mice were recorded. The first three rows represent the data of three nodes shown in (b,c,e). (g) The statistics of the two forms of microglia-node contact shown in (f). (h) Diagram showing the definition of microglial primary and secondary processes. (i) The statistics of nodal contacts performed by different microglial components in the case of intermittent (top) and constant contact (bottom). (j) Percentage of nodes with microglial contact over time. The grey area indicates a small region where locates all the data points. Scale bars, 20 μm in (b) and (e), 10 μm in (c), 5 μm in (d).

We further investigate how microglia performed continuing and intermittent contact differently. Given that NR can be contacted by different microglial components, including soma, primary processes, and secondary processes (Fig. 1h), we characterized the events accordingly (Fig. 1i). In our analysis of 18 cases of continuing contact, we found that the majority (15 out of 18) were initiated by microglial soma or primary processes. Conversely, in the case of intermittent contact, the vast majority of nodes (48 out of 49) were contacted by the secondary processes. These findings are consistent with the notion that microglial soma and primary processes exhibit less dynamism compared to the secondary processes under normal physiological conditions. Consequently, once the microglia-node contact is established with its soma or primary processes, it is prone to be statically maintained for at least several hours. We noted that when examining the microglial contact with individual nodes, it is difficult to ascertain a pattern governing the interaction. However, upon analysing all nodes collectively, the percentage of nodes with microglial contact remained relatively stable over time, with 55% to 70% of NR being contacted by microglia at any given time point. (Fig. 1j). Additionally, all NR were found to have contact with microglia within a three-hour timeframe. These findings support the notion that microglia play a crucial role in closely monitoring axons at NR, highlighting their essential contribution to the maintenance of normal axonal function.

### Localized wrapping contact of microglia with NR in response to axonal injury

In our previous work, we observed a more frequent contact at the proximal nodes after laser injury^19^. Particularly, in some cases, microglial processes were intensively recruited to wrap the nodes. Expanding on our prior research, we conducted a detailed investigation to determine whether the observed wrapping contact by microglia occurred exclusively at the node nearest to the site of injury or extended to multiple nodes along the injured axon. In the mouse spinal cord, the typical distance between two adjacent NRs ranges from 200 μm to 2 mm, and this distance can vary significantly depending on the physiological properties of the axons^45^. As a result, our current imaging methods have a limited field of view (FOV), allowing us to image only two adjacent nodes together. To perform highly localized laser axotomies, we targeted a distance >200 μm distal from the two nodes of interest. The distance was determined to avoid the chemotactic effect of laser injury interfering with the behaviour of microglia surrounding the nodes, as discussed previously^19^. Time-lapse imaging with a 5-10 min time interval was started 0.5 hr before the laser axotomy, and the imaging continued to track the axon-microglia dynamics following the injury for an hour. Our findings revealed a striking pattern: exclusively the nodes in immediate proximity to the lesion sites exhibited microglial wrapping contact, while the second-closest nodes demonstrated intermittent contact, closely resembling the pattern observed under physiological conditions (Fig. S1b, movie S2). Subsequently, we sought to investigate whether wrapping contact could also occur at distal nodes by inducing laser injury between two nodes. Remarkably, even in cases where there was disconnection from the neuronal cell body, we consistently observed microglial wrapping contacts at the distal nodes (Fig. S1c). In all five cases studied, both the closest distal and proximal nodes exhibited wrapping by microglial processes (Fig. S1d), indicating that these wrapping contacts at nodes are a localized response independent of neuronal soma participation. Notably, microglia processes tended to wrap the NRs first and then gradually wrap paranodal and even juxtaparanodal areas towards the lesion site after axonal injury.

Next, we conducted laser axotomy in close proximity to NR (50-100 μm) to explore whether the injury-induced chemotaxis could disrupt microglial wrapping behaviour. We consistently observed wrapping contact at NR in the majority of cases (4 out of 5) (Fig. S2a) despite the attraction of microglial processes towards the adenosine 5’-triphosphate (ATP) or adenosine 5’-diphosphate (ADP) released from the lesion site^6^. Thus, microglial wrapping response to axonal injury at NR is prominent and little affected by the chemotactic interference from the lesion site. We then randomly determined the lesion site at a 50-300 μm distance distal from the targeted nodes with no other NR exists in between (Fig. 2a). Not surprisingly, wrapping contacts were observed in the vast majority of nodes (20 out of 21, Fig. 2b) within 1 hour post injury (hpi), and interestingly, the response time of this wrapping reaction had no obvious correlation with the axotomy-NR distance (Fig. S2b). The wrapping contact between microglia and axons, although rapidly established within an hour, is short-term and lasts only for a few hours. At 2 hpi, approximately 7.7% of the axons lost microglial wrapping at their nodes, and this percentage increased to 15.4% at 4 hpi (Fig. 2c). By 24 hpi, only one wrapping contact was observed out of 13 observations, while all other wrapping contacts were either completely lost or transformed into normal transient contacts (Fig. 2a, c).

**Fig. 2.**
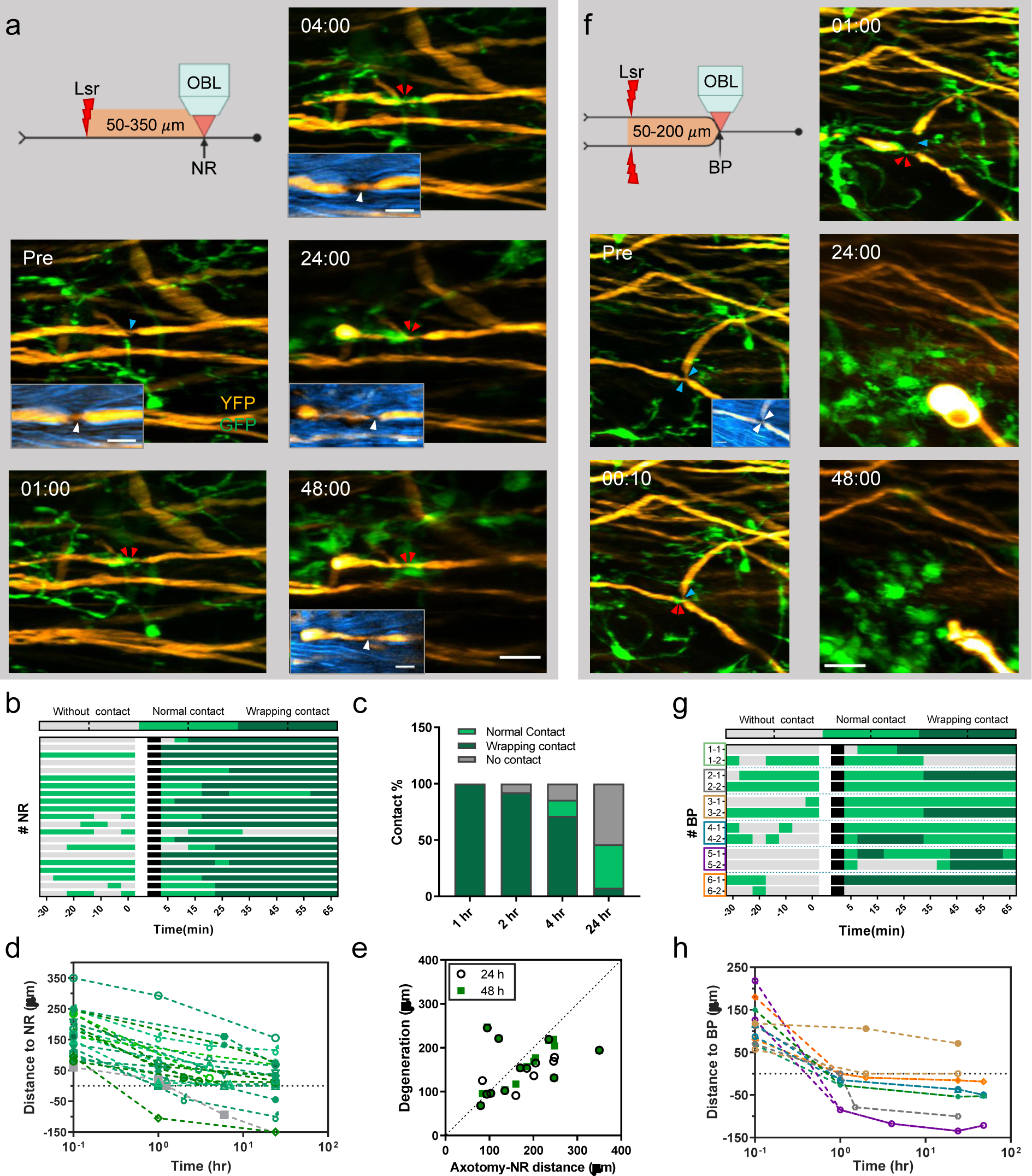
Microglial protection of axons from acute degeneration through physical wrapping at NR. (a) Illustration of the experimental design and the corresponding time-lapse imaging results of the microglia-node interaction before and after laser axotomy. Blue arrowheads indicate nodes without microglial contact and red arrowheads indicate nodes contacted by microglia (green). Double red arrowheads indicate wrapping contact at NR. Time post injury is presented as hr: min. Insets, overlay of myelin SRS image (blue) and axon YFP image (orange). White arrowheads indicate the location of NR. Images in (a) are representative of 15 axons observed in 9 mice. (b) Heatmap showing microglia-node contacts from 0.5 hr before to 1 hr after laser axotomy. The blank area indicates the interruption of time-lapse imaging when laser injury was performed. Time-lapse imaging sessions were restarted 5 minutes after the injury. A total of 21 nodes from 9 mice were recorded. (c) Change in wrapping contact events over time. Only wrapping contacts occurring within 1 hour post-injury (hpi) were considered in the statistics. (d) The distance between the proximal ends of injured axons and the proximal NR over the first 24 hr after injury. The grey colour indicates the data of axons without wrapping contact. Curves across the dashed line indicate the axonal degeneration events with nodal breach. (e) Plot of degeneration distance against the distance between the axotomy site and NR. Data points above the dashed 45° line represents the incidence of NR breach. Colocalization of the circle and square data points indicates the cessation of axonal degeneration from 24 to 48 hpi. A total of 16 axons were recorded. (f) Illustration of the experimental design and the corresponding time-lapse imaging results of the microglia-node interaction before and after double branch axotomy. Time post injury is presented as hr: min. Insets, overlay of myelin SRS image (blue) and axon YFP image (orange). White arrowheads indicate the location of NR. (g) Heatmap showing microglia-node contacts from 0.5 hr before to 1 hr of two daughter branches after double branch axotomy. A total of 6 axons with secondary branch points (BPs) from 4 mice were recorded. The color of the box of the row title corresponds to the color of the curve in (h). (h) The distance between the proximal ends of injured axons and BPs over the first 24-48 hr after injury. The same color indicates the daughter branches belonging to the same BP. Scale bars, 20 μm in (a), (f); 10 μm for all the insets in (a) and (f).

### Protective role of microglial wrapping contact at NR against acute axonal degeneration

Following axotomy, injured axons undergo bidirectional degeneration, with the proximal ends retrogradely dying back in a fragmentation-dominant manner, and the distal segments undergoing Wallerian degeneration^21,24^. In this study, we focused on the retrograde degeneration of the injured axon. Based on the previous evidence that most axon loss occurs within 4 hrs after a lesion and the proximal ends become stable after about 30 hr^21,24^, we tracked injured axons for ∼4 hrs on day 0 and reimaged them in the following two days. Our results demonstrate that in most cases (17 out of 20), regardless of the axotomy-NR distance, the injured axons ceased their acute degeneration slightly before the NR, leaving a small segment without breaking the nodes before 4 hpi (Fig. 2a, d). Consistent with previous results^21,24^, the bursts of degeneration typically happened in the first few hours and became subsided within 4 hr after axotomy (Fig. 2d). In the following two days, axons that stopped before nodes on day 0 usually shortened slightly more but still left their proximal ends before the NR (13 out of 16) (Fig. 2a, e). Also, these injured axons stopped their demyelination at NR within the first two days (Fig. 2a). Considering prior research indicating that secondary branch points (BPs) where the entering spinal axons first bifurcate can function as barriers against retrograde acute degeneration when one branch is damaged while the other remains intact^24^, we conducted a separate analysis of secondary BPs. Both the immunostaining and *in vivo* imaging results show that the node at BP also bifurcates and both daughter branches have paranodal structures at BP (Fig. 1d (ii) and Fig. S3), which is different from the normal nodes.

In the case of secondary BPs, we initially induced damage to one of the daughter branches at a distance of 50-300 *μ*m from the BP. Microglial wrapping contacts occurred in all cases (4 axons), and all daughter branches stopped degeneration before the BPs by day 2. Similarly, the demyelination also stopped at the BP with the other branch kept intact (Fig. S3). Next, we injured both of the two daughter branches sequentially. In response to this double-branch axotomy, microglia wrapped only one branch of the BPs in most cases (5 out of 6), and the wrapped branch was not consistently the first to be injured (Fig. 2f-g, movie S3). It should be noted that the ablation point should not surpass the subsequent node (Fig. S4). By contrast, within 2dpi, most axons (5 out of 6) underwent degeneration beyond the BP (Fig. 2h). Collectively, except the double-branch axotomy case, most axons stopped their acute degeneration and demyelination before/at nodes whatever the nodal structure is.

Our findings imply that microglial wrapping behaviour functions as a mechanism to safeguard NR, preventing axonal degeneration from compromising its integrity. To test this hypothesis, we aimed to completely abolish microglia-node contact by ablating microglia without affecting the survival of other CNS cells. After three-weeks treatment of PLX3397 (290 mg/kg), which pharmacologically inhibits microglial pro-survival receptor CSF1R^46,47^, microglia in the spinal dorsal column were effectively eliminated by approximately 82% (Fig. 3a-b). To ensure complete elimination of microglial contact both before and after laser axotomy, nodes located far away from the remaining few microglial cells within the FOV were selected for the study, despite the presence of a few microglial cells (Fig. 3c-h). In the normal node case, 75% (9 out of 12) axons degenerated out of NR within 4 hr after axotomy (Fig. 3c, f). A few axons (8.3%, 1 out of 12) further degenerated to break the nodes in the next 20 hrs and almost all the tracked axons showed little change from 24 to 48 hpi (Fig. 3f). For branch points with collateral sprouting/branching, injury of the main shaft resulted in similar rapid axonal degeneration without stopping before/at branch point, leading to the elimination of the other uninjured small branch in all 3 cases (Fig. S5). However, in the case of secondary BP, injury of only one daughter branch didn’t lead to the breach of BPs in all 6 cases (Fig. 3d). These BP structures exhibited stiffness that resisted axonal degeneration, likely due to the presence of the other intact branches, which could counteract the loss of wrapping contact. Not surprisingly, when all the daughter branches were damaged, they degenerated to breach the BP with a significantly high probability similar to the control group (5 out of 6) (Fig. 3e, h). In conclusion, we found that microglia formed specialized wrapping contact at NR which arrested axonal proximal ends before or at NR. This protective mechanism prevents long-distance axonal degeneration during the acute phase. However, this protective effect seems to play an inferior role in regulating the fate of the first bifurcating BP after injury of the daughter axons.

**Fig. 3.**
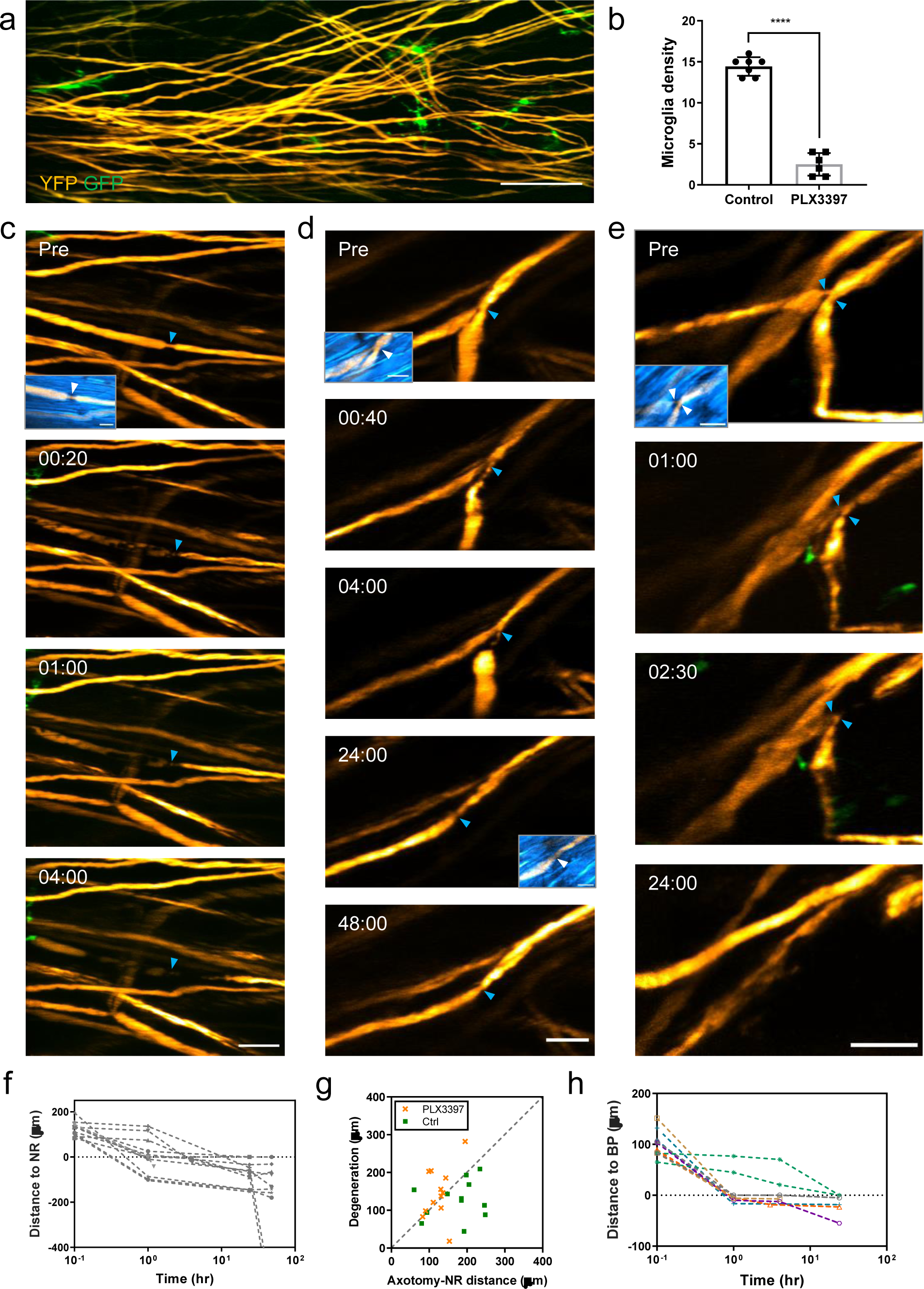
Acute axonal degeneration in the absence of microglia-node interaction. (a) An *in vivo* spinal cord image shows significantly decreased microglial density after three-week treatment of PLX3397 (290 mg/kg). Orange, YFP labelled axons; Green, GFP labelled microglia. Scale bar, 50 μm. (b) Statistics of microglial density before (n = 7) and after three-week treatment of PLX3397 (n = 6). Error bar, standard deviation (SD). Unpaired two-tailed *t*-test: ****P < 0.0001. (c-e) In the absence of microglial contacts with nodes, axonal response to laser axotomy performed proximal to a normal NR (c), on a daughter branch of a secondary branch point (BP) (d) and on both daughter branches of a secondary BP (e). Time post injury is presented as hr: min. Scale bar, 20 μm. Insets, overlay of myelin SRS image (blue) and axon YFP image. White arrowheads indicate the position of the targeted normal nodes and branching nodes. Scale bar, 10 μm. (f) Distance between proximal ends and NR of the injured axons over the first 24 hr after injury. A total of 12 axons from 6 mice were recorded. (g) Plot of degeneration length against the distance between the axotomy site and NR at 4 hpi. Data points above the dashed line represent the incidence of nodal breach. Axonal degeneration data shown in Fig. 2d was reprocessed as the ‘Ctrl’ datasets here. (h) Distance between proximal ends and secondary BPs of the injured axons over the first 24 hr after injury. The same color indicates the daughter branches belonging to the same BP. A total of 6 axons from 5 mice were recorded.

### Modulation of wrapping contact by microglial P2Y12 receptors via purinergic signalling following axonal injury

Our study have demonstrated microglia-axon interactions at NR in both physiological and pathological conditions. To unravel the molecular mechanism underlying the communication between microglia and nodes, our study centered on investigating the P2Y12 receptor because this receptor holds exclusive expression in microglia within the CNS and is distributed across their membranes, including the delicate fine processes^16,18,48,49^. Specifically, purinergic signalling through microglial P2Y12 receptor has been shown to mediate similar microglial processes recruitments to neuronal soma or dendrites after acute brain injury^16^ or excessive neuron activation^50–52^. In addition, P2Y12 receptor function also modulates physiological neuron-microglia interaction at somatic junctions^16^. Therefore, we investigated the crucial role of microglial P2Y12 receptor signalling in mediating microglia-axon interactions at NR. In the first experimental study, a highly selective and potent P2Y12 receptor inhibitor PSB0739 (PSB) was administered by intrathecal injection. The optimal treatment dose in the spinal cord was determined by assessing the microglial response to laser ablation. It was found that the injection of a 10 μl solution containing 0.1 mg/ml of PSB significantly delayed the chemotaxis of microglial processes evoked by laser injury compared to the control (no treatment) and vehicle control (saline injection) group, though it did not result in the complete abolition of this response (Fig. S6a-b). Following the treatment, microglia exhibited a minor reduction in ramification compared to their pre-treatment state (Fig. S6c), aligning with previous findings^53^. We also observed that the PSB effect decreased with time but still showed significant inhibition at 6 hr after treatment (Fig. S6a), consistent with the study in brain^16^. In order to mitigate the potential impact of PSB’s diminishing inhibition at later time points, all imaging of microglia-node interaction was conducted within 6 hours following PSB injection. To exclude the potential effects caused by intrathecal injection, we also analysed microglia-node interaction in the presence of vehicle (saline) injection. The results showed that the both the physiological interaction between microglia and NR and microglial wrapping response following axonal injury were not affected by the vehicle injection (Fig. S7). Next, to assess the influence of P2Y12 signalling on physiological microglia-node interaction, a 1-hour time-lapse imaging of the same region was performed before and 30 minutes after PSB treatment (Fig. S8a). Microglial contacts at each node before PSB injection was used as the baseline for future comparison. In our experiments, we did not find that physiological microglia-node contacts were altered by inhibiting microglial P2Y12 receptor, both in terms of total contact time and the maximum contact duration (Fig. S8b-f, movie S4). The percentage of contacted nodes at each time point also showed little change after PSB treatment, consistent with previous results obtained from myelinated cerebellar slices^18^.

Next, we examined whether P2Y12 receptor was involved in the nodal wrapping contact in response to axonal injury. The *in vivo* imaging sessions commenced 30 minutes after the administration of PSB and were conducted for a duration of less than 6 hours. At each node, cellular dynamics were continually tracked for 1.5 hours, including 0.5 hr before and 1 hour after laser axotomy. Extra snapshots were later captured with 1-2 hr time intervals to record the acute axonal degeneration progress. We found that wrapping contacts were almost absent after injury with PSB treatment. Out of 21 tracked normal nodes, only two normal nodes showed temporary wrapping contact, which did not prevent axonal degeneration from breaching the nodes (Fig. 4a-d). In the case of axons lacking wrapping contacts, acute degeneration typically did not halt either before or at normal nodes in the majority of instances (83.3%, 10 out of 12). With the wrapping contacts largely abolished by blocking P2Y12 receptor with PSB, axons showed similar acute degeneration processes as in the microglial-depletion case. Hence, although the inhibition of P2Y12 signalling has no effect on the physiological microglia-axon contact at NR, P2Y12 receptors plays a crucial role in triggering the microglial wrapping response at nodes to safeguard injured axons during the acute degeneration stage.

**Fig. 4.**
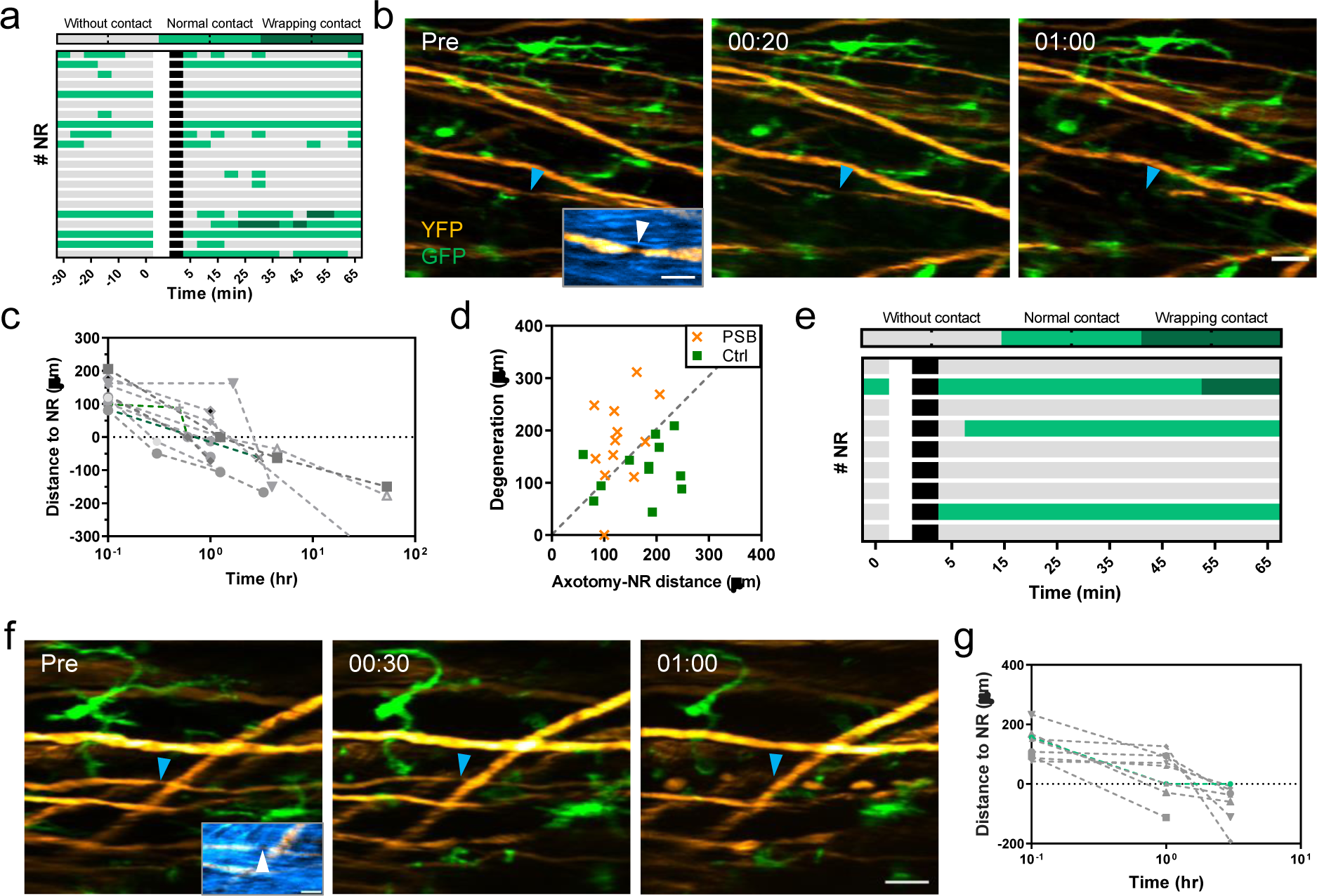
Purinergic signaling is involved in the microglia-node wrapping contact following axonal injury. (a) Heatmap showing microglia-node contacts from 0.5 hr before to 1 hr after laser axotomy in the PSB0739 (PSB) treatment group. The blank area indicates the interruption of time-lapse imaging when laser injury was performed. Time-lapse imaging sessions were continued 5 minutes after the injury. A total of 21 nodes from 11 mice were recorded. (b) Time-lapse images at indicated time points showing the microglia-axon dynamics and interactions in response to axonal injury. Time post injury is presented as hr: min. Blue arrowheads indicate the absence of microglia-axon contact at NR. Insets, overlay of myelin SRS image (blue) and axon YFP image (orange). White arrowheads indicate the location of NR. (c) Distance between the proximal ends and NR of injured axons over the first 24 hr after injury in the PSB group. The gray color indicates the data of axons without wrapping contact. Curves across the dashed line indicate the axonal degeneration events with nodal breach. (d) Plot of degeneration distance against the distance between the axotomy site and NR at 4 hpi. Data points above the dashed line represent the incidence of NR breach. Axonal degeneration data shown in Fig. 2d was reprocessed as the ‘Ctrl’ datasets here. (e) Heatmap showing microglia-node contacts from 5 mins before to 1 hr after laser axotomy in the NPPB treatment group. The blank area indicates the interruption of time-lapse imaging when laser injury was performed. Time-lapse imaging sessions were continued 5 minutes after the injury. A total of 9 nodes from 4 mice were recorded. (f) Time-lapse images at indicated time points showing the microglia-axon dynamics and interactions in response to axonal injury. Time post injury is presented as hr: min. Insets, overlay of myelin SRS image (blue) and axon YFP image (orange). White arrowheads indicate the location of NR. (g) Distance between the proximal ends and NR of injured axons over the first 3 hr after injury in the NPPB group. The gray color indicates the data of axons without wrapping contact. Curves across the dashed line indicate the axonal degeneration events with nodal breach. Scale bars, 20 μm in (b) and (f); 10 μm for all the insets in (b) and (f).

P2Y12 is a chemoreceptor for ADP/ATP^53,54^ and its inhibition impairs the ADP/ATP-induced chemotaxis of microglial processes^6,48,53,55^. It is known that ATP can be directly released by axons through the VAAC activated by microscopic axonal swelling^56–58^. During action potential firing, axonal swelling was observed together with ATP release^57^, which subsequently attracted microglial processes to the periaxonal area^58^. In this study, we also observed morphological change in axons, especially at NRs, after laser ablation (Fig. 2a, Fig. S1b-d and S2a). Hence, it is plausible that the nodes have the capability to release ATP through VAAC, thereby attracting microglial processes to establish wrapping contact after injury. To explore this hypothesis, our initial approach involved the direct observation of ATP release at NR *in vivo*. We achieved this by attaching a fluorescent ATP sensor to the axonal membrane through virus injection^59^. Unfortunately, the attempts to directly visualize ATP release at NR by swelling axons were unsuccessful. This was probably due to the low expression level of the sensor in myelinated axons and the limited sensitivity of the method (Fig. S9). Then we used a non-selective VAAC inhibitor, 5-nitro-2-(3phenylpropylamino) benzonic acid (NPPB)^58^, to address whether the VAAC contributes to the microglia-node interaction after injury. Similar to PSB, VAAC blocker affects microglial ramification and impairs its laser response^60,61^. Injection of 10 ul 1.5mg/ml NPPB significantly abolished laser injury-evoked chemotaxis of microglial processes within 1.5-hour post injection (Fig. S10a). After treatment, 88.9% (8 out of 9) of injured axons were not wrapped by microglia and degenerated out of nodes within 3 hours (Fig. 4d-g).

It must be emphasized that though the impact of PSB and NPPB on microglia following laser injury was comparable (Fig. S6a and S10a), their mechanisms were expected to differ. While PSB impeded the chemotaxis by inhibiting P2Y12 receptor in microglia, NPPB was expected to reduce the ATP release through VAAC after injury, without affecting microglial response to ADP/ATP because a previous study demonstrated that VAAC in microglia did not contribute to P2Y12-dependent chemotaxis^61^. To further confirm that NPPB indeed affects ATP release in mouse spinal cord, we used the aforementioned ATP sensor^59^ to monitor ATP release in the spinal cord *in vivo*. Corresponding AAV virus were injected into the spinal cord to label the dorsal column axons, which were imaged three-week later under the treatment of saline, PSB and NPPB. As a result, in the control group treated with saline, the ATP sensor showed weak basal fluorescence signal, which were boosted immediately after a local laser injury. This observation was consistent with previous findings^59,62^. In the PSB group, the ATP sensor was lightened by laser ablation as in the control group, which indicates PSB has no effect on ATP release. By contrast, NPPB treatment reduced the response of ATP sensor markedly after laser injury within 0.5h post injection and the effect decreased slightly at 1.5h post injection (Fig. S10b). Hence, it was confirmed that NPPB effectively reduced ATP release, consequently impacting microglial wrapping behaviour in response to axonal injury, while PSB inhibited the P2Y12 receptor. However, it remains unclear whether NPPB exerts an indirect or direct effect on the P2Y12 receptor simultaneously. Nevertheless, taken together, purinergic signalling plays a crucial role in regulating the wrapping contact between microglia and NRs following injury.

### Involvement of nodal sodium channels in modulating microglia-node interaction

Myelinated axons have clustered ion channels at NR, which allows the action potential regeneration and saltatory conduction. Previous studies have shown that neuronal activities regulate neuron-microglia interaction behaviour^16,50,52,63^. In particular, an ex vivo study of Purkinje neuron in organotypic cerebellar slices proved that physiological microglia-node contacts were affected by the change of neuronal activity and dependent on the potassium level within the nodal area^18^. Therefore, we next explored whether the neuronal activities similarly modulate microglia-node contact in spinal dorsal column. Since mice were anesthetized with isoflurane during *in vivo* imaging, the occurrence of spontaneous neuronal activities was reduced. To activate the dorsal root ganglion (DRG) neurons as well as its central branch in the spinal cord, we performed electrical paw stimulation for mice, which has been a common method to increase sensory neuronal activities ^64–68^. By inserting two needles into mice footpad, we delivered constant bipolar currents to initiate electrical impulses, enabling activation of the downstream dorsal horn neurons^44,65,69^.

In the following *in vivo* experiments, 1hr time-lapse imaging were performed before and after 10-min electrical stimulation to visualize the dynamic microglia-node contact in the physiological circumstance (Fig. 5a-b, movie S5). After electrical stimulation, the percentage of contacted NRs (37 NRs in total) was significantly increased but no wrapping contact occurred (Fig. 5c-d). The results indicated the involvement of neuronal activities in modulating physiological microglia-node communication. However, wrapping contacts cannot be triggered by increasing neuron activities in the physiological condition. Next, we examined the influence of ion channel functions related to neuron activity on microglia-node contact following axonal injury. We first focus on the voltage-gated sodium channels (NaV), which is responsible for the rising phase of action potential ^70,71^. Tetradotoxin (TTX), a NaV inhibitor, was intrathecally administered to inhibit neuronal activities during *in vivo* imaging. We found that application of TTX doesn’t affect microglial morphology and motility as well as its response to laser injury^18^ (Fig. S11). However, it alters microglial wrapping behaviour as well as axonal acute degeneration pattern. Blocking the sodium channel with TTX, 56.25% (9 out of 16) of axons were not wrapped by microglia following injury. Surprisingly, for axons without wrapping contact, only 37.5% (3 out of 8) of them degenerated out of NRs within 4 hours post injury (Fig. 5e-g). This degeneration rate was much lower compared to those treated with PSB, NPPB and PLX3397 (over 83%). As a result, voltage-gated sodium channels were involved in the regulation of microglia-node contacts after axonal injury. Blocking these channels can prevent further axonal degeneration, even in axons without protective wrapping contact by microglia.

**Fig. 5.**
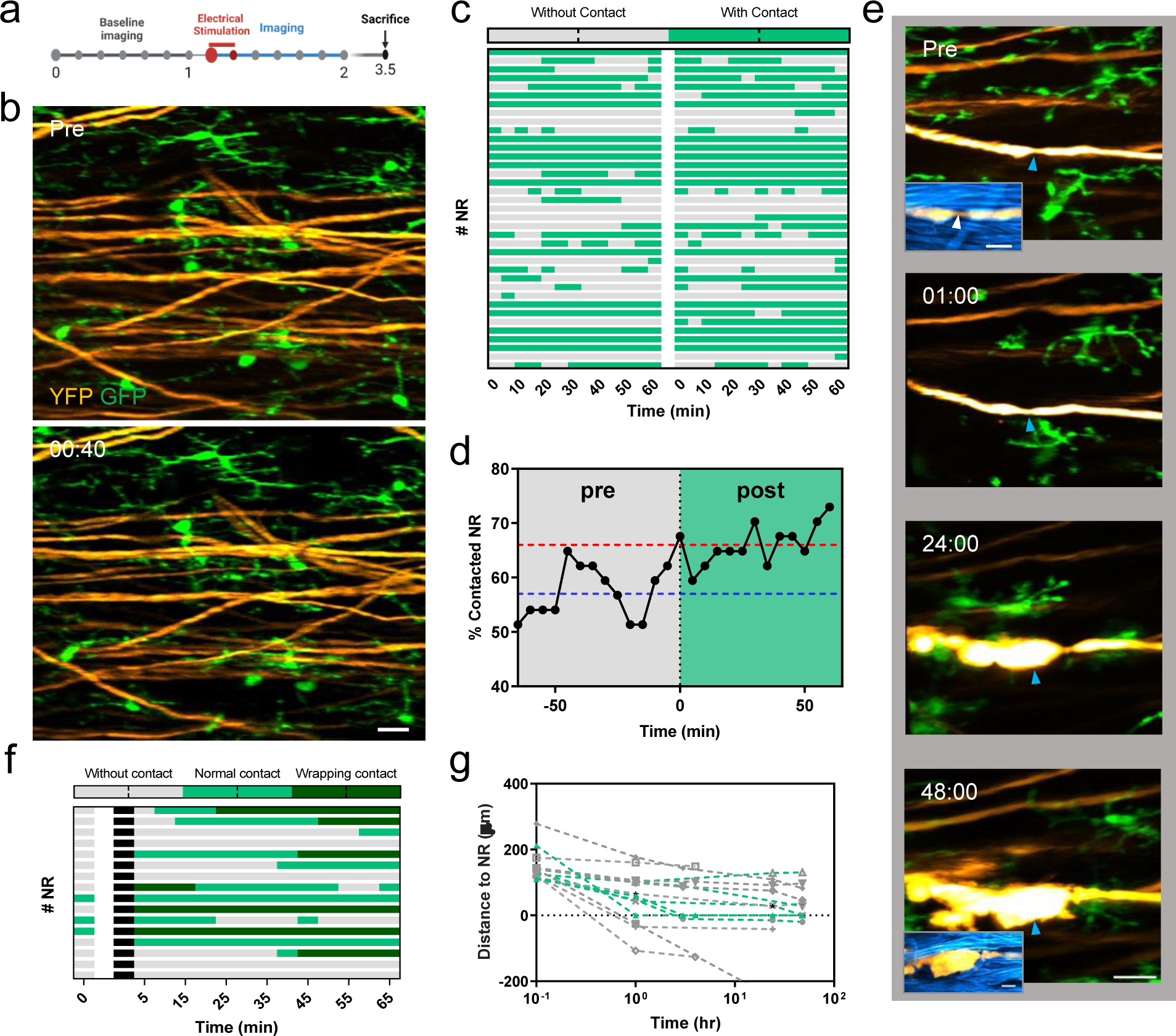
**Regulation of microglia-node interaction by neuronal voltage-gated sodium channels in physiological and pathological conditions. (**a) Illustration of the experimental design. (b) Representative images of microglia (green) and axons (orange) before and 40 mins after electrical stimulation. Scale bar: 20 μm. (c) Heatmap showing microglia-node contacts during 1-h time-lapse imaging before and after electrical stimulation. The blank area indicates the time interval for stimulation. Images were taken every 5 minutes. A total of 37 nodes from 3 mice were recorded. (d) The percentage of contacted NR by microglia in the 37 recorded nodes before and after electrical stimulation. The grey vertical dashed line indicates the start point of electrical stimulation. The purple and red dashed line indicate the mean percentage value of contacted NR before (∼57%) and after (∼66%) stimulation, respectively. (e) Time-lapse images at indicated time points showing the microglia-axon dynamics and interactions in response to axonal injury in the TTX treatment group. Scale bar, 20 μm. Insets, overlay of myelin SRS image (blue) and axon YFP image (orange). White arrowheads indicate the location of NR. Scale bar, 10 μm. The axon did not degenerate out of node within 48 hpi. Time post injury is presented as hr: min. (f) Heatmap showing microglia-node contacts from 5 mins before to 1 hr after laser axotomy in the TTX group. The blank area indicates the interruption of time-lapse imaging when laser injury was performed. Time-lapse imaging sessions were continued 5 minutes after the injury. A total of 16 nodes from 8 mice were recorded. (g) Distance between the proximal ends and NR of injured axons over the first 48 hrs after injury in the TTX group. The gray color indicates the data of axons without wrapping contact. Curves across the dashed line indicate the axonal degeneration events with nodal breach.

Considering the modulating effect of potassium release at nodes on microglia-node interaction in cultured slices^18^, we sought to determine if this mechanism also operates *in vivo* within the spinal cord for microglia-node contact. In the experiment, a high concentration of tetraethylammonium (TEA), a broad-spectrum inhibitor of potassium channels, was applied through intrathecal injection to effectively suppress the outward potassium current. Successful TEA treatment was confirmed by the initiation of mouse muscle trembling. Microglia-node contacts at the same site were continually imaged before and after TEA treatment. Contrary to the previous results obtained from the brain slice, physiological microglia-node contact dynamics and percentage of contacted NRs was not changed significantly by TEA treatment (Fig. S12a-b). Based on these findings, it can be concluded that the blocked potassium release at nodes by TEA does not play a role in the physiological microglia-node interaction in the spinal dorsal column. The observed variation in results is likely attributed to the differences in imaging preparation methods and the specific population of targeted neurons. Additionally, in our study, 93.7% (15 out of 16) of axons exhibited wrapping by microglia following injury (Fig. S12c-d). Furthermore, the degeneration rate of normal nodes (28.5%, 4 out of 14) was comparable to that of the control group (Fig. S12e). These findings indicate that the inhibition of potassium channels by TEA does not have an impact on microglia wrapping contact at nodes after laser axotomy.

It should be noted that while TEA can widely block the 6TM family of potassium channels, it works poorly for the two-pore domain potassium channels^72–74^, which exist at the surface of both NR and microglia^75,76^. Therefore, the possibility of potassium participation in the microglia-node interaction cannot be simply ruled out due to its insensitiveness to the TEA treatment. Subsequently, we employed a two-pore domain K+ channel blocker to further explore the role of potassium in microglia-node communication. Here tetrapentylammonium (TPA) was selected to inhibit the THIK-1 channel, which is mainly expressed in microglia as one of the highly expressed two-pore domain K(+) channels. THIK-1 activity controls microglial membrane potential, which further regulates its ramification and surveillance but doesn’t affect its directed motility^76^. Therefore, inhibition of the THIK-1 channel with TPA should not affect the function of microglial P2Y12 receptor. In our experiment, TPA was delivered intrathecally and half-an-hour time-lapse imaging was performed before and after its treatment. As a result of TPA application, microglia showed a significantly decreased ramification (Fig. S13a) but similar response to laser injury as in the control condition (Fig. S13b), which is consistent with previous results and confirms TPA having taken effect during the *in vivo* imaging. After axonal injury, microglia still showed prominent wrapping response by extending its processes to the nodes, little affected by TPA treatment (Fig. S13c). Taken together, the microglial wrapping contact at nodes after axonal injury was not affected by the potassium fluxes.

## Discussion

As microglia cells are the primary immune cells in the CNS, their morphology is often used as an *in vivo* indicator of inflammatory activity. The absence of morphological alterations, such as process retraction and decreased ramification, is considered minimally invasive and indicative of non-activated microglia^77–81^. In this study, we utilized a multimodal microscopy system in conjunction with an optically-cleared intervertebral window to enable non-invasive *in vivo* imaging of the spinal cord. This approach allows us to visualize the near-native cellular dynamics and interactions over time without triggering activation of microglial morphology. By employing this technique, we were able to identify the specificity and functionality of microglia-axon communication at NR with minimal interference, providing valuable insights into their interactions in the CNS. We demonstrated that, under physiological condition in normal spinal cord, microglia established physical contacts with axons at NR in a preferred random scanning manner, while maintaining contact with a certain percentage of nodes at all times. In all observed cases, the dynamic intermittent contacts were established by microglial processes rather than their soma. It is worth noting that the movement of microglial soma is considerably slower, by several orders of magnitude^53,82^. Similar dynamic intermittent contacts have also been observed between microglial processes and neuronal soma^16^ as well as dendrites^83^. Our study provides further evidence for the hypothesis that the substantial motive of maintaining microglial process constant movement is to establish direct membrane-membrane contacts^10^.

Axonal injury serves as a prominent pathological feature in various conditions, including spinal injuries and other inflammatory-mediated neurological disorders. While the role of microglia in maintaining homeostasis is well-established, understanding their response to axonal injury and their contribution to subsequent pathological processes is an area of research that has garnered considerable attention. Identifying microglial function during axonal degeneration as well as the following regeneration trials can potentially offer new therapeutic targets to develop effective treatment strategies. Following spinal cord injury, microglia become activated and display various phenotypes in response to inflammation, showing neurotoxic or neuroprotective effects. While previously observed interactions between activated microglia/macrophage and injured axons usually lead to phagocytosis^38,84^ or secondary axonal degeneration^36,85^, our results offer a new perspective on the function of physical interactions between microglia and injured axons. By performing localized axonal injury at the targeted position using laser axotomy, we observed a highly-responsive microglial wrapping contact at NR of the injured axon. After the establishment of wrapping contacts, it is observed that acute/subacute axon degeneration is more likely to halt at or before the nodes. However, in the absence of microglial wrapping response, this protective effect is no longer observed. Therefore, microglial processes converging at NR act as a barrier for axonal acute degeneration, indicating a neuroprotective role of the physical contact between microglia and damaged axons. Furthermore, this specific neuroprotective function of microglia brings to mind a previous study that demonstrated how axonal branch points, characterized by a surviving intact branch, also serve as barriers for preventing axonal degeneration^24^. Given that branch points within NR also attract microglial process wrapping following injury, we hypothesize that the barrier role of branch points in preventing axonal degeneration may be attributed to microglial contacts in a similar manner. Hence, we performed laser axotomy at one and both branches of a branch point with microglial contact abolished. In the case of branch points formed by the central branch first bifurcating after entering the spinal cord (secondary branches), damaging only one branch usually leads to the total elimination of the injured branch but not breaching the branch points with or without wrapping contact (Fig. 3d, S3). In contrast, when all branches are simultaneously damaged, there is a significant increase in the rate of breaching the branch point, regardless of whether wrapping contact occurs or not (Fig. 2g-h, 3h). Therefore, the surviving branch plays a dominant role in stabilizing the branching structure after injury, while microglial wrapping contacts at the branch points contribute little to determine the axonal fate. In addition to exploring the response of branch points formed by secondary bifurcation, we also investigated the response of other branching forms, such as collateral sprouting/branching. These branch points with a small axonal branch/sprout emerging from the main shaft exhibited a similar performance to the normal NR in response to the main shaft injury (Fig. S5). With elimination of microglial contact, the injured main shafts usually went through quick retrograde degeneration without stopping when encountering the branch point. Regarding collateral branches, it is still unclear whether the observed differences from secondary bifurcation points are due to a potential hierarchical response to injury based on the order in which the branches are affected^30^.

In this study, we revealed that microglial processes were recruited to NR of injured axons in a P2Y12-dependent manner, which effectively blocks the retrograde axonal degeneration from further progression when reaching the NR. Both the application of PSB0739 (P2Y12 antagonist) and NPPB (VAAC inhibitor) similarly impaired the microglia ramification and response to laser injury, abolishing microglial wrapping contact with nodes. Although we were unable to directly observe ATP release by the axon at the nodes, the inhibition of ATP release at the lesion site by NPPB indirectly confirmed the involvement of ATP in this microglia-node interaction (Fig. S10b). However, we can’t exclude the possibility that it is ATP released by other cells around the nodes that contribute to microglial wrapping response, as NPPB also blocks the VAAC channels in other none-neuron cells in the spinal cord^6,57,86–88^. Besides, it remains unknown whether NPPB impairs P2Y12 activity of microglia indirectly by reducing microglia ramification^48^, which further affects the wrapping contact between nodes and microglia. Future studies may investigate the source of ATP release in the vicinity of the nodes where wrapping contact occurs.

Although our study demonstrated that electrical stimulation effectively enhanced neuronal activity, resulting in an increased percentage of NRs being contacted by microglia in the normal spinal cord, it should be noted that the exact proportion of YFP-labeled neurons that are activated by electrical stimulation remains undetermined, even though the electrical currents were applied to the entire hind limb foot pad^44,69^. Moreover, after axonal injury, the application of TTX to some extent prevented microglial wrapping responses. These *in vivo* results are in line with a prior investigation that showed a significant decrease in microglia-node contacts following TTX treatment in organotypic cerebellar slices^18^. Furthermore, a previous study indicated that nodal potassium fluxes also contribute to modulating microglia-node interactions^18^. Both TEA and TPA were found to inhibit microglial interactions with nodes. However, in our study, the inhibition of potassium channels using TEA did not affect microglia-node contacts in the normal spinal cord. Additionally, both TEA and TPA showed minimal effects on wrapping contacts following axonal injury. This inconsistency is likely attributed to different experimental conditions in the respective investigations. Although TEA and TPA didn’t have an effect on regulating microglia-node interaction, the role of potassium flux in the process remains elusive. It is unclear whether TREK-1 or TRAAK, which are the thermosensitive and mechanosensitive two-pore-domain potassium (K_2P_) channels responsible for rapid action potential repolarization at the NRs, rather than voltage-gated K(+) channel^70^, are more likely to be the principal potassium release channel at nodes. It needs further study to drive a conclusion.

Interestingly, besides inhibiting microglial wrapping response, TTX treatment also prevents axonal degeneration from breaching the node in the absence of wrapping contact. Over the past few decades, progresses have been made in understanding the molecular mechanisms underlying axonal degeneration. It has been shown that axonal degeneration is typically initiated by an increase in intra-axonal calcium concentration, followed by the activation of calpain, a calcium-dependent protease^21,89–92^. Reduced calcium influx by inhibiting calpain can attenuate progressive damage of axon after SCI^21,40^. Previous studies also demonstrated that blockage of voltage-gated sodium channel by TTX could ameliorate tissue loss and block the onset of acute white matter pathology after SCI^93,94^, which may contribute to impaired calcium influx or reduced inflammatory responses after TTX treatment^95–97^. Additionally, the mechanism behind how microglial wrapping contacts at the NR prevent axonal acute degeneration is yet to be explored. In addition to calcium influx during axonal degeneration, mitochondrial dysfunction has also been reported as an early ultrastructural sign of axonal damage, which precedes axonal swelling and degeneration^20,90^. Collectively, these findings suggest that microglia may protect axons from further degeneration by impeding extracellular calcium influx as the way TTX works or regulating mitochondrial activity near the NR, which requires further investigation in future studies.

After demonstrating that microglia establish neuroprotective contacts with damaged axons during the acute phase of degeneration, the subsequent question to address is whether specialized physical contact also persists in the subacute and chronic phases, potentially contributing to axonal regeneration attempts. While previous studies have observed physical interactions between activated microglia and proximal ends of damaged axons in the brain and spinal cord^38,98^, few studies have examined their role in axonal regeneration, largely due to the limited regeneration capability of the adult CNS. However, evidence from studies of the developing brain has shown that microglia can actively participate in the refinement of neuronal connections by promoting and guiding the outgrowth of axonal tracts^99,100^. Therefore, it would be interesting to investigate whether microglia retain the capability to promote and guide the regeneration of damaged axons in the adult CNS with enhanced regeneration capability achieved by advanced genetic modulation or a preconditioning stimulus^101,102^.

Lastly, it is important to emphasize that the spinal cord injury model employed in this study involved localized laser axotomy, which enabled the investigation of microglia-neuron interactions in a well-controlled manner. While our results demonstrate the neuroprotective function of microglial wrapping contacts at NR in stabilizing injured axon structures, they also offer novel insights for the development of effective therapeutic strategies for axonal injury and neurological disorders. However, it is important to note that the microenvironment at the site of laser injury differs from that in established animal models of large-scale SCI, including sharp transection, compression injury, crush injury, or complete transection. Therefore, it should be approached with caution when attempting to generalize the findings of this study regarding microglial behavior to the context of large-scale injuries. Further research is necessary to develop models that accurately mimic clinical conditions of spinal cord injury and enable minimally invasive imaging, in order to gain a comprehensive understanding of microglial behavior in such contexts.

## Methods and Material

### Animals and surgery

Cx3Cr1^GFP/+^(B6.129P2(Cg)-Cx3cr1tm1Litt/J)^103^ mice with microglia labelled with EGFP were used to examine microglial dynamics under various pharmacological treatment in the spinal cord. Thy1-YFP/Cx3Cr1^GFP/+^ transgenic mice were generated by crossing Cx3Cr1-GFP with Thy1-YFP (Tg(Thy1-YFP)HJrs/J)^104^ transgenic line for *in vivo* study of microglia-axon interaction. The mice were kept in groups of two to four in a controlled environment with access to food and water. The environment was maintained at a temperature of 22-25°C with 40-60% humidity, and a standard 12-hour light-dark cycle was followed. . Experiments were conducted using adult mice of either sex, aged 10 to 20 weeks. All animal procedures in this work were conducted in accordance with the guidelines of the Laboratory Animal Facility of the Hong Kong University of Science and Technology (HKUST) and were approved by the Animal Ethics Committee of HKUST.

The surgical procedures were carried out on mice that were under anaesthesia and placed on a heating plate at 37°C, unless otherwise stated. The anaesthesia was induced by injecting a mixture of ketamine (80 mg/kg) and xylazine (10 mg/kg) into the peritoneal cavity. All surgical instruments were sterilized either by autoclaving or disinfecting them with 70% ethanol. The surgical area was made sterile by covering the benchtop with clean drapes. After surgery, the mice were given a subcutaneous injection of 1 mg/kg atipamezole to help them recover from the anaesthesia. Post-operative pain was managed by administering 0.1 mg/kg buprenorphine through subcutaneous injection.

### Intervertebral window

To prepare the optically-cleared intervertebral window for *in vivo* imaging, a modified version of a previous protocol was followed^19^. Briefly, a small incision was made over the T11-T13 vertebra to expose the dorsal tissue after hair removal and skin disinfection with iodine. Muscle tissues and tendons above and alongside the vertebra were removed, which was then held in place by two clamping bars on a custom-designed stabilization stage. Next, muscle tissues and tendons in the cleft between T12-T13 were gently removed, leaving ligamentum flavum intact. After exposing the spinal cord with LF lying over, care should be taken to avoid touching the window surface during the surgery. Sterile saline was used to flush away excess blood, and keep the surgical area clean. Sterile gauze pads were used to control bleeding if necessary. Once the spinal cord was fully exposed with the ligamentum flavum lying over it, a coverslip was placed on the clamping bars with the 0.5-1 mm interspace between the spinal cord and cover slip. This space was then filled with Iodixanol (60% w/v, D1556, Sigma-Aldrich) for *in vivo* imaging. To visualize microvascular vessels, mice received a retro-orbital intravenous injection of 50 μl Texas Red dextran (70 kDa, 1 mg/100 μl in saline, Invitrogen) before *in vivo* imaging. After imaging, Iodixanol was removed and the wound was cleaned with saline and gauze pads. To protect the intervertebral window, liquid Kwik-Sil (World Precision Instruments) was applied and left to cure for approximately three minutes. The skin was then sutured to close the wound and burn cream (Betadine) was applied on the sewed skin to prevent infection. Mice were placed on a heating pad until fully recovered from anesthesia and were housed individually thereafter. For reimaging through the same intervertebral window, the sutured skin was reopened and the topical Kwik-Sil was removed. Any newly generated tissues adhering to the vertebra were detached if necessary to ensure stable clamping of the vertebra. Granulation tissues on the surgical site were carefully peeled off to expose the LF. Optical clearing was then applied as previously described.

### Intrathecal administration

Most drugs were diluted with sterile saline to achieve an appropriate final concentration for injection. Specifically, the dose used for each drug was 0.1mg/ml for PSB0739 (Tocris), 0.5 ug/kg body weight for TTX, 1mg/ml for TEA (Sigma) and 15mg/ml for TPA-Cl (sc-251210, Santa Cruz Biotechnology). For intrathecal administration of NPPB (MedChemExpress, MCE), drugs were dissolved and diluted with a new solvent formed by adding 10% DMSO, 40% PEG300, 5% Tween-80 and 45% saline in sequence, following the protocol provided by the MCE. The final used dose of NPPB was 1.5 mg/ml. The final dose of different drugs were dependent on the response of microglia or physical reaction of mice.

To perform intrathecal injection, after shaving the back fur and disinfecting the skin with iodine, the skin overlying the vertebral column was incised. The injections were performed by inserting a 31G needle connected to a 300 ul syringe between L5-L6 vertebrae of the anesthetized mice. Tail flick were observed as the sign for successful entry of the needle in the intradural space. ∼10 μL of targeted solution were slowly injected and the needle were retained for approximate 1 min after injection.

### Virus injection

To label the spinal cord axons with GRAB ^59^, the T12-T13 intervertebral window was exposed and AAV2/9-hsyn-cATP1.0 (2.00E+12vg/mL, BrainVTA) was injected into the spinal cord through the window using a micro injector attached to a fine glass pipette. The injection site was 0.4 mm from the midline on both sides of the spinal cord, and the depth was 0.2 mm below the dura. Totally 500 nL AAV was injected at each point at a speed of 100 nL/min.

### Microglia depletion

For microglia depletion, mice were fed PLX3397 (MedChemExpresss) formulated in AIN-76A standard chow (290mg/kg) for a continuous period of three weeks.

### Integrated SRS and TPEF microscopy

The multimodal non-linear optical microscope system used in this study is similar to previous work^19,44,105^. An integrated optical parametric oscillator (OPO, picoEmerald S, APE) connected to an 80 MHz mode-locked Ytterbium fiber laser was used as the laser source for SRS imaging. The OPO output includes a tunable pump beam (780-960 nm) and a Stokes beam of 1031 nm with 2 ps pulse duration. The Stokes beam is modulated at 20 MHz by a built-in electro optical modulator for high-frequency phase-sensitive SRS detection. A tunable femtosecond Ti:sapphire laser (Chameleon Ultra II, Coherent) served as the laser source to excite two-photon fluorescence (TPEF). The ps and fs beam were first collimated and magnified by a pair of lenses to match the size a pair of Galvo XY-scanning mirror (3 mm, 6215H, Cambridge Technology). Before being directed to the scanning mirror, ps and fs beams were combined by a polarizing beamsplitter (PBS) (CCM1-PBS252/M, Thorlabs) with a half-wave plate (SAHWP05M-1700, Thorlabs) to adjust the polarization of fs beam. A scan lens (SL50-CLS2, Thorlabs) and a tube lens (TTL200-S8, Thorlabs) was placed after the scanning mirror to conjugate it with the rear pupil of the objective lens (XLPLN25XSVMP2, 25×/1.05 NA, Olympus).

For SRS imaging, the pump beam was tuned at 796 nm to visualize the myelin sheath at 2863.5 cm^-1^, corresponding to its Raman peak at C-H region. The backscattered pump signal was directed to a large area silicon photodiode (10 mm × 10 mm, APE) using another PBS (CCM1-PBS252/M, Thorlabs). A dichroic short-pass filter (69–206, short-pass at 700 nm, Edmund) was positioned following the PBS to separate the SRS signal from the fluorescence signal for detection. Before the photodiode, a short-pass filter (86–108, short-pass at 975 nm OD4, Edmund) and a band-pass filter (FF01-850/310, Semrock) were installed to completely eliminate the Stokes beam. A high-speed lock-in amplifier (LIA) was employed to demodulate the pump signal received from photodiode, allowing highly sensitive detection of stimulated Raman loss.

For two-photon imaging, the femtosecond laser set at 920 nm was used to excite the fluorescence signals from YFP, GFP, ATP sensor, and Texas Red. The TPEF signal collected by the objective was reflected to the photo detection unit using a long-pass dichroic mirror (FF665-Di02, Semrock). An interchangeable dichroic beam splitter (FF488-Di01-25 × 36 or FF518-Di01-25 × 36, Semrock) was placed after the dichroic shortpass filter to differentiate the fluorescence with difference wavelength before detection. Two filter sets were placed before two current photomultiplier (PMT) modules (H11461-03 and H11461-01, Hamamatsu) to reject the laser beam and isolate the desired fluorescence signal. One filter set (FF01-715/SP-25, Semrock; FF01-525/50, Semrock or HQ620/60X, Chroma) was utilized for YFP/Texas red signal detection, the other filter set (FF01-720/SP25, Semrock; 67-029, 488-512 nm band pass, Edmund) was used for GFP/ATP sensor signal detection. The two PMT outputs were directed into two current amplifiers for signal conversion and amplification. The output of current amplifiers and LIA was digitized by a data acquisition device (PCIe-6363, National Instrument) for image reconstruction. A custom C# program was developed to control all the hardware components for image acquisition.

### In vivo imaging

*In vivo* imaging typically commenced 1 hr post-surgery, unless stated otherwise. After surgical preparation, mouse was fixed onto a stabilization stage^19^ which was then placed on a five axis stage, enabling three-axis translation and ±15∘ pitch and roll flexure motion. The holding plate was lowered to slightly suspend the mouse’s body, allowing extra space for chest movement and minimizing the motion artifacts from breathing. If needed, the mouse’s head was held steady by two head bars to further reduce motion artifacts. A heating pad set at 37℃ was positioned beneath the mouse’s body to maintain warmth during imaging. Throughout the imaging session, the stage’s roll and pitch angles were adjusted to ensure the imaged region perpendicular to the optical axis as possible. A 10 × objective (NA = 0.45, Nikon) was generally used to acquire a large FOV image for region of interest (ROI) selection and navigation. A 25× water immersion objective ((XLPLN25XSVMP2, Olympus) was used to acquire high-resolution two-photon and SRS imaging of the targeted region. For two-photon imaging, the post-objective power varied from 40 to 60 mW and 30 to 50 mW for 10× and 25× objective, respectively, depending on the transparency of the window and cellular fluorescence intensity. For SRS imaging of myelin sheath, the power of the pump and Stokes beam was set to be 50–70 mW and 60–80 mW for 25× objective, respectively. During *in vivo* imaging, Iodixanol had to be replaced every 1-2 hours to maintain optimal clearing effects. Imaging was usually performed 5-10 mins after Iodixanol application when the optical clearing starts to function effectively. Isoflurane (0.5-1%) was used to keep mice under light anesthesia.

For electrical stimulation during *in vivo* imaging, a previous protocol was used with minor revisions^44^. Briefly, two 27-gauge hypodermic needles were inserted into the hind limb foot pad to deliver the electrical current from a custom-made bipolar constant current stimulator. A sponge was placed under the hind limb to stabilize the piercing needles. The stimulation was applied with 0.28 mA constant current amplitude for 10 minutes and confirmed by onset of perceptible hindlimb twitching.

### Laser axotomy

Laser axotomy is performed with high-power femtosecond laser pulses under the guidance of two-photon fluorescence imaging. In practice, the 920 fs beam was parked at the targeted position in the just-captured the TPEF image for 1-4 s, with an average power of 200-300 mW. Laser axotomy is based on a multiphoton ionization and thereafter plasma-mediated ablation process. New fluorescence can be generated during the process, which can be used to mark the lesion site and evaluate the size of lesion.

### Immunohistology

Mice were anesthetized by intraperitoneal injection of K.X.S at a lethal dose and then perfused with PBS followed by 4% (w/v) PFA (Sigma-Aldrich). The spinal cord was dissected and immersed in 4% PFA at 4℃ overnight. The fixed spinal cord samples were dehydrated in 15% (w/v) for 12 h and then in 30% sucrose overnight. Dehydrated samples were cut into 3 mm segments, and embedded into OCT compound (Tissue-Tek) at -80℃ overnight. Then the sample was cut into 30 *μm* coronal sections using CryoStar NX70 Cryostat (Thermo Scientific). Tissue sections were blocked with 4% NGS containing 0.1% Triton X-100 for 1 hour then incubated with primary antibody (1:500 diluted by 4% NGS) overnight. Samples were washed 3 times with PBS followed by incubating with the secondary antibody (1:500 diluted by 4% NGS) for 2 hours. Thereafter, samples were washed 3 times with PBS before being mounted on the slides with anti-quench mounting reagent. Fluorescent images were taken by ZEISS Axio Scan Z1 Slide Scanner. Antibodies used in this study were: Rabbit anti-Caspr (Ab34151, Abcam), Goat anti-rabbit IgG (H+L) Alexa Fluor™ 555 (A21429, Invitrogen), and Goat anti-rabbit IgG (H+L) Cyanine5 (A10523, Invitrogen).

### Image processing and analysis

To compensate the motion artifacts caused by breathing and heartbeats, image registration for microglia and axon images was performed using a previous method^19,44^. Since microglia and axon images were captured simultaneously by two detection channels, they experienced the same distortion and required identical transformation matrix for correction. In brief, each detection channel captured 10 frames (512 × 512 pixels) at a 2-Hz frame rate for every depth. Intra-frame registration was first applied to one channel by registering each frame to their averaged image at each depth using the ‘bUnwarpJ’ plugin^106^ in imageJ^107^. The elastic transformation parameters for each registration were saved and then applied to the other channel’s images. The final slice at each depth was the average of the registered frames, resulting in minimal motion artifacts and enhanced signal-to-noise ratio. The ‘MultiStackReg’ plugin was used for registration between slices at different depths of one channel and the transformation information was saved and later applied to the other channel. After reconstructing 3D image stacks for different time points, the 2-channel stacks were merged, converted to hyperstacks and aligned using the YFP signal of axons with the ‘Correct 3D drift’ plugin. All registration procedures were conducted using the Fiji macro programming language. Large-FOV images (Fig. S4 and Fig. 3a) were obtained by stitching sub-images with the ‘Pairwise stitching’ plugin^108^. Unless otherwise specified, all fluorescence images shown in the figures are maximal intensity projection of an image stack, while SRS images are displayed as single image slice.

To calculate microglial density in the spinal cord, microglial image stack of a 300 *μ*m*300 *μ*m*45 *μ*m imaging volume were projected as maximal intensity projection (MIP) and image contrast was adjusted to saturate 0.3% pixels. Microglial cells with intact soma in the imaging volume were counted for density quantification. The cell density presented for each mouse was the average value of 3-6 data points from different regions.

For quantification of microglial morphology, cells with clear and intact morphology were chosen in an unbiased manner from each FOV. The analysis was performed using Fiji and MATLAB (The MathWorks) scripts based on the published methods^76,109^. In short, microglial MIP images were adjusted to saturate 1% of pixels and then smoothed. The adjusted images were thresholded and performed Hessian matrix enhancement^110,111^ to obtain the optimized binary image with minimal process loss using a home-built GUI in MATLAB. To determine the ramification index, the MATLAB functions ‘bwarea’ and ‘bwperim’ (8-connected neighborhood) were applied to the binary images, yielding area and perimeter values. The ramification index is calculated as:

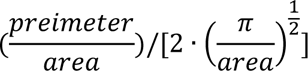

The quantification of microglial response to laser ablation followed a previous method with minor modifications^6^. After laser ablation, time-lapse microglial image stacks were capture, maximally projected and smoothed before adjusting the contrast to saturate 1% of pixels. These images were then manually thresholded to create binary images of microglia for further analysis. Since laser ablation produced a small autofluorescence sphere of around 10 *μ*m in diameter, the center of this sphere can be regarded as the centroid of both inner area X (35 *μ*m in radius) and outer area Y (70 *μ*m in radius). The number of positive pixels in area X (R*_x_*(t)) over time were was and compared to the initial image taken immediately after the ablation (R*_x_*(0)). To account for variation in microglia number in the outer area Y across different experiments, microglial responses at different time points can be determined by:

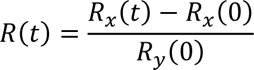

For microglia-nodes contact analysis, the location of NRs on the axons were validated by merging the myelin SRS and axon TPEF image. Next, axon images were merged with microglia images. Contacts were recognized as the 3D overlap of the fluorescence of microglia and NRs. Wrapping contact was defined when the NRs were completely surrounded by microglial fluorescence.

For axon degeneration analysis, the distance between the lesion site and the NRs were recorded. In addition, the distance of proximal terminal of the axons to the NRs at different time points was monitored based on the YFP signal of axons.

### Statistical Analysis

Statistical analysis and data visualization were performed using GraphPad Prism 7 software. All measurements were performed on more than 3 mice. The data are presented as mean ± SD, or ± SEM and α =0.05 for all analyses. Data normality was first checked using the Shapiro-Wilk normality test. Normally distributed data were analyzed using paired, unpaired two-tailed t-test and two-way ANOVA test. Non-normally distributed data were analyzed using paired or unpaired non-parametric t-test (Wilcoxon test or Mann-Whitney test). No statistical methods were used to determine the sample size.

## Acknowledgments

This work was supported by the Hong Kong Research Grants Council through grants 16102123, 16103215, 16148816, 16102518, 16102920, T13-607/12R, T13-605/18W, C6002-17GF, C6001-19E, N_HKUST603/19 and the Innovation and Technology Commission (ITCPD/17-9), and the Area of Excellence Scheme of the University Grants Committee (AoE/M-604/16, AOE/M-09/12) and the Hong Kong University of Science & Technology (HKUST) through grant 30 for 30 Research Initiative Scheme.

We thank the histology and microscopy core of Biosciences Central Research Facility, HKUST (Clear Water Bay).

## Author contributions

W.W, Y.H and J.Y.Q conceived the research idea and designed the experiments; W.W built the imaging systems; Y.H and W.W performed animal surgery and imaging experiments; Y.C performed the IT injection, virus injection and histology study under the supervision of K. L; Y.H and W.W analyzed the data with the help of Y.F and S.H; W.W and Y.H wrote the paper with inputs from all other authors.

