## Supplementary Info for "Microglial-Mediated Prevention of Axonal Degeneration in the Injured Spinal Cord: Insights from an *In Vivo* Imaging Study"

### Supplementary Figures

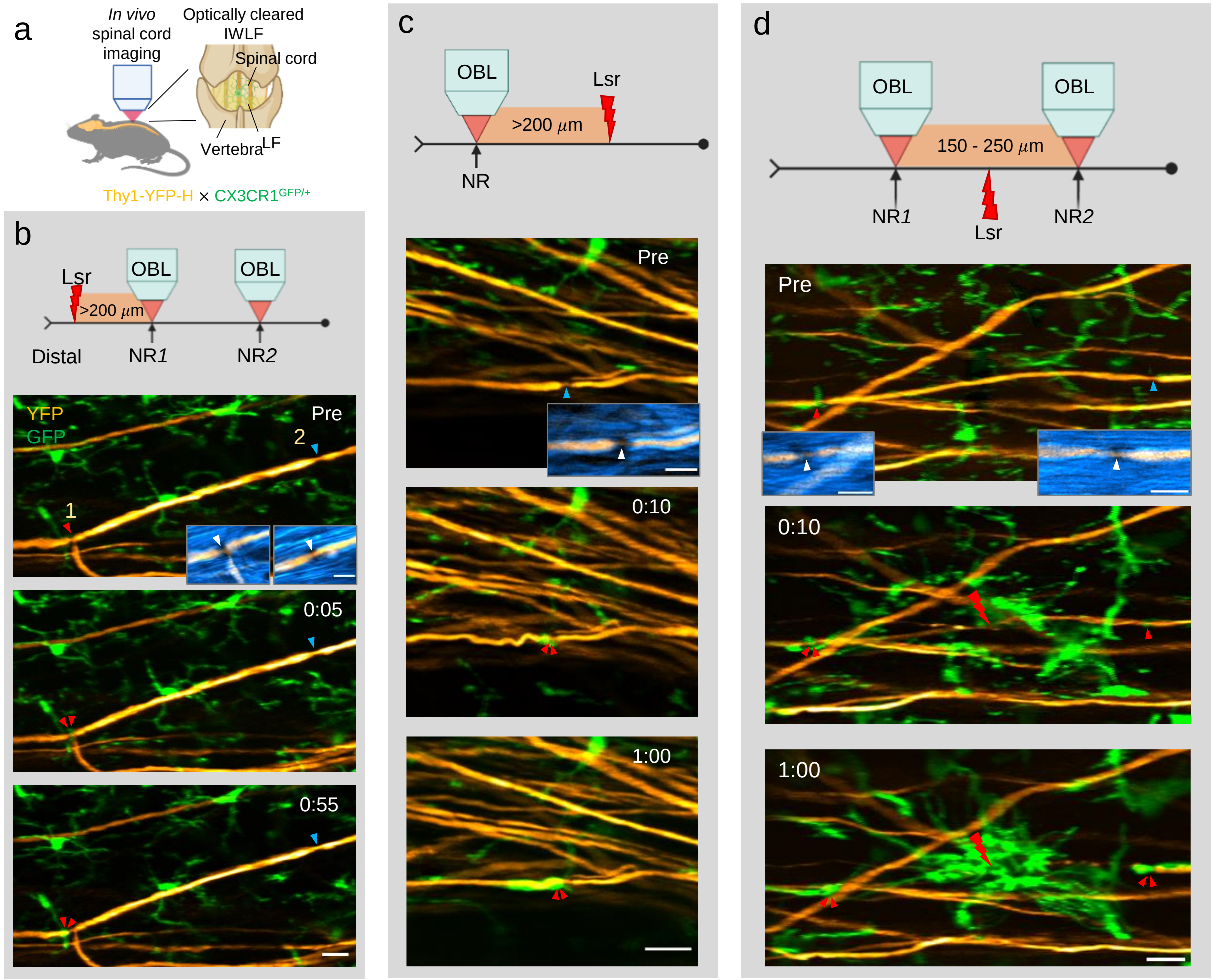


**Fig. S1 Microglial processes wrap the closest node of Ranvier (NR) of injured axons.** (a) Schematic of *in vivo* imaging of mouse spinal cord through an optically cleared intervertebral window with ligamentum flavum (IWLF). (b-d) Illustration of the experimental design and the corresponding time-lapse imaging results of the microglia-node interaction before and after laser axotomy. Blue arrowheads indicate nodes without microglial contact and red arrowheads indicate nodes contacted by microglia (green). Double red arrowheads indicate wrapping contact at NR. Time post injury is presented as hr: min. Insets, the overlay of the myelin SRS image (blue) and axon YFP image (orange). White arrowheads indicate the location of NR. Images in (b-d) were representative of 5 axons. Scale bars, 20 µm in (b-d), 10 µm in all the insets.

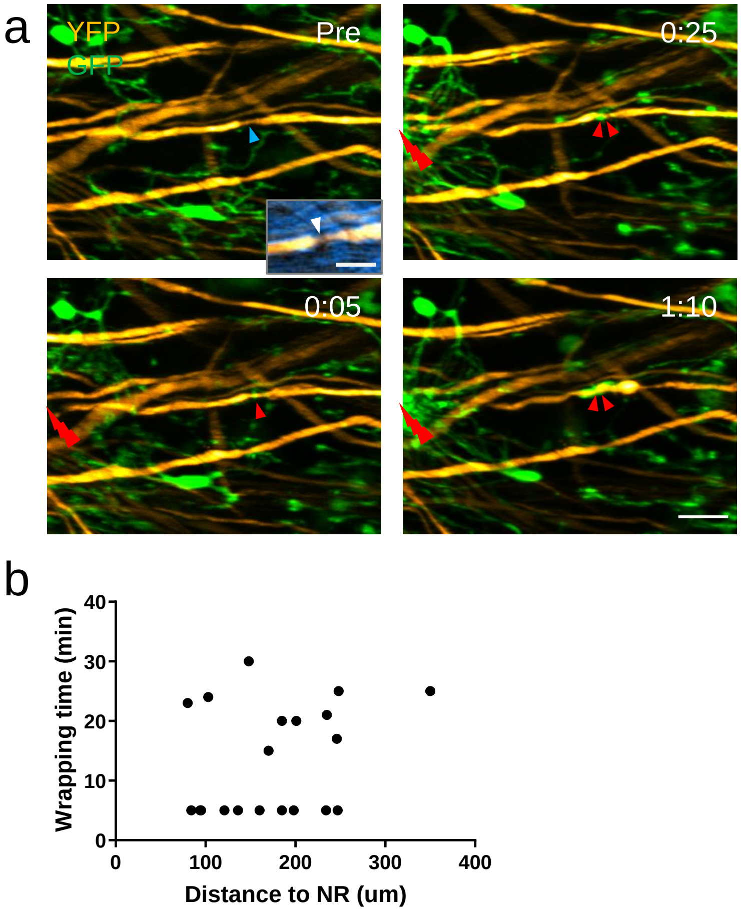


**Fig. S2 Distance of lesion site to NRs does not affect wrapping contact between microglia and NRs.** (a) Representative images for the microglia-node interaction after a short distance injury (~80 μm). Blue arrowheads indicate nodes without microglial contact and red arrowheads indicate nodes contacted by microglia (green). Orange, YFP-labeled axons. Double red arrowheads indicate wrapping contact at NR. Red lightning bolts indicate the ablation points. Time post injury is presented as hr: min. Scale bar, 20 μm. Inset, the overlay of the myelin SRS image (blue) and axon YFP image (orange). White arrowhead indicates the location of NR. Scale bar, 10 μm in the inset. (b) Plots of the correlation between the wrapping time and the distance of ablation to NR. The Pearson coefficient (r) is 0.2688.

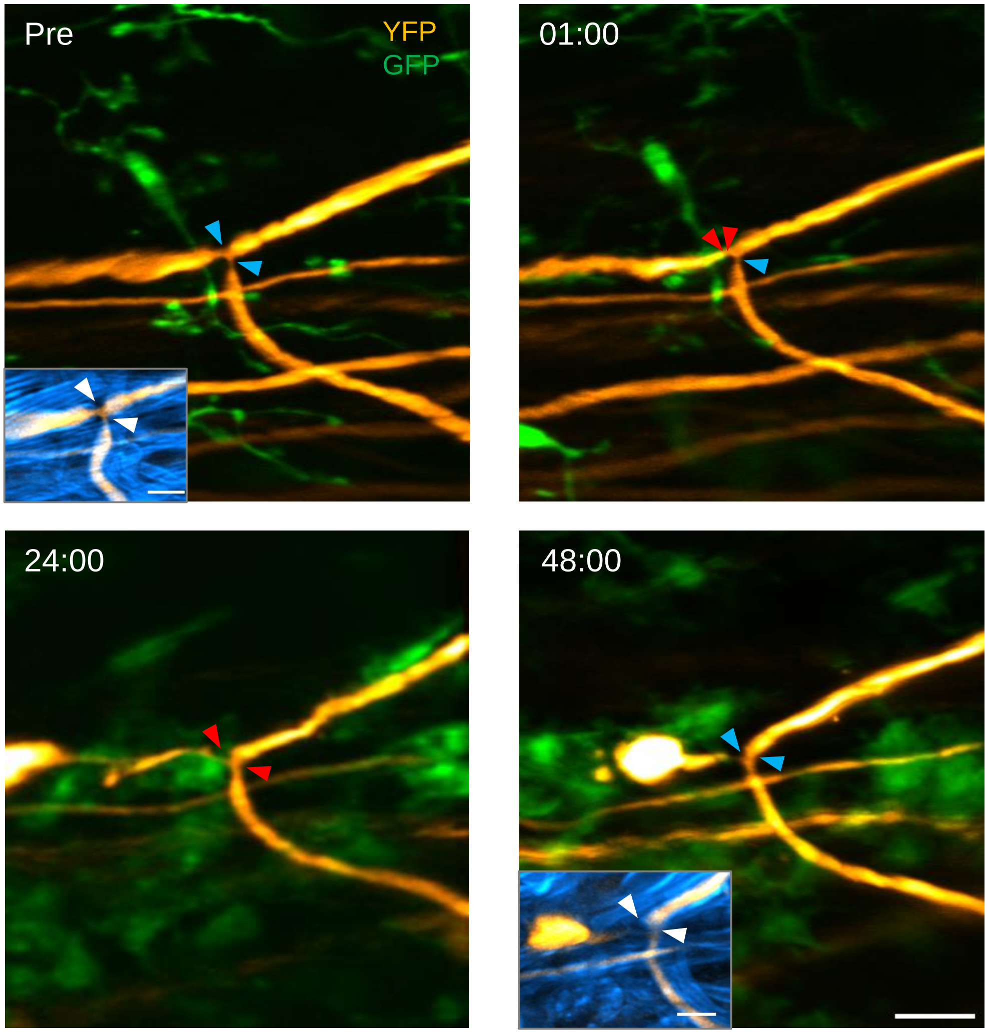


**Fig. S3 Microglia-axon dynamics in response to single daughter branch axotomy.** Time-lapse imaging of microglia-axon dynamics at a secondary BP before and after single daughter branch injury. Blue arrowheads indicate nodes without microglial contact and red arrowheads indicate nodes contacted by microglia. Double red arrowheads indicate wrapping contact at NR. Insets show the location of NR and BP (white arrowheads). Orange, YFP-labelled axons, green, GFP labelled microglia, blue, the myelin SRS. Scale bars, 20 µm, 10 µm for all the insets. Time is represented as hr: min.

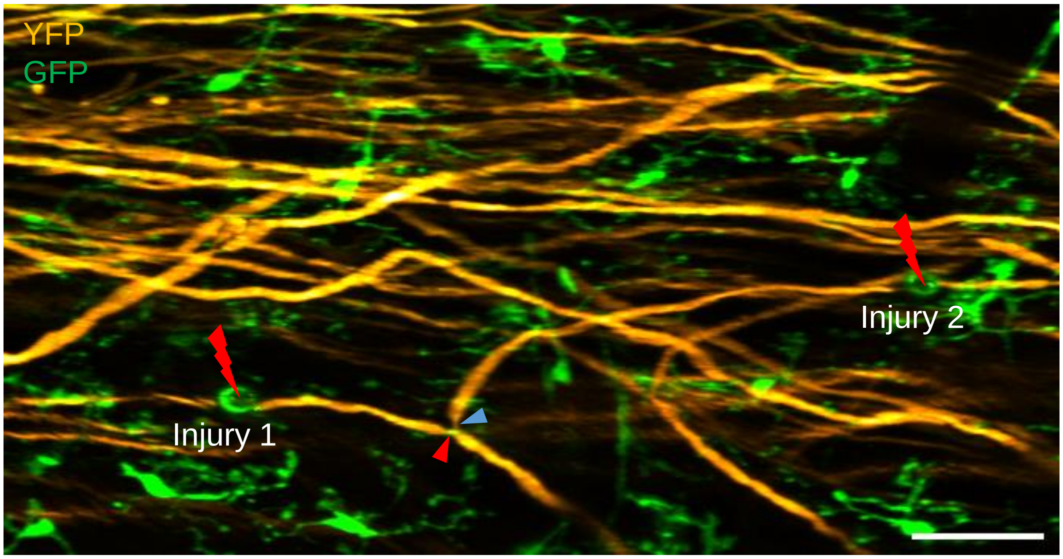


**Fig. S4 Double branch axotomy of an axon bifurcating at the secondary branch point.** Orange, YFP-labelled axons; green, GFP-labelled microglia. Red lightning bolts indicate the injury sites. Arrowheads indicate the position of branch point. Scale bar, 50 μm.

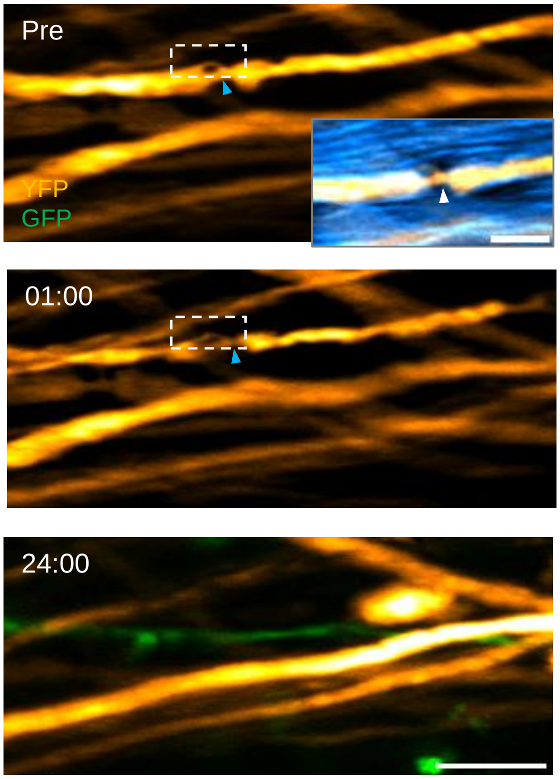


**Fig. S5 Degeneration of an injured axon with collateral branch/sprout in the absence of microglial contact after microglia depletion.** Time-lapse images at the indicated time points showing the degeneration of axons (orange) after injury. Green, GFP-labeled microglia. White dotted boxes show the collateral branch. Axon degenerated out of the branch points in the absence of microglial contact. Insets, overlay of myelin SRS image (blue) and axon YFP image (orange). White arrowheads indicate the location of NR. Scale bars, 20 μm, 10 μm for the inset. Time is represented as hr: min.

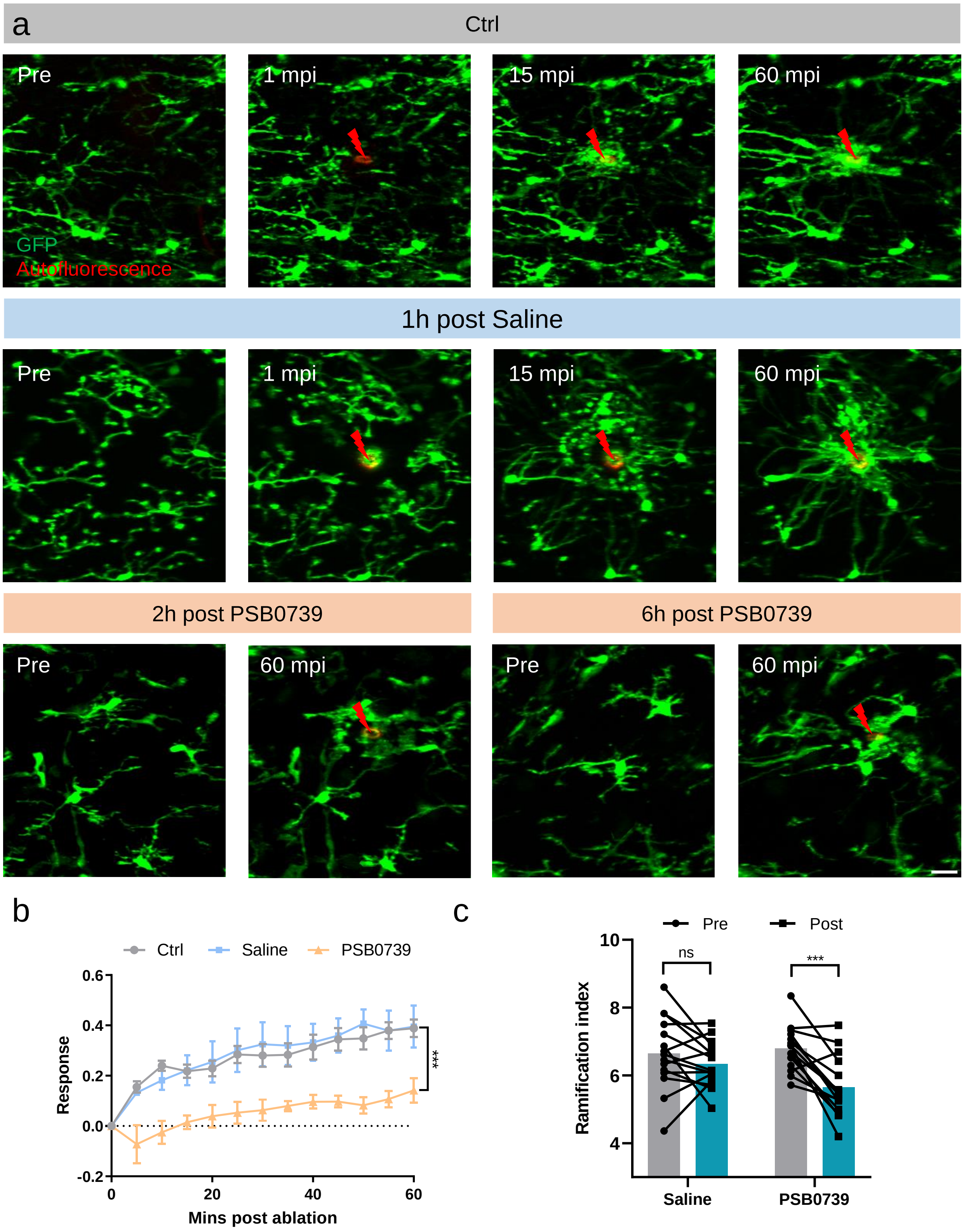
a
Ctrl
Pre
1 mpi
15 mpi
60 mpi
GFP
Autofluorescence
1h post Saline
Pre
1 mpi
15 mpi
60 mpi
2h post PSB0739
6h post PSB0739
Pre
60 mpi
Pre
60 mpi
b
Ctrl
Saline
PSB0739
Response
Mins post ablation
***
c
Pre
Post
ns
***
Ramification index
Saline
PSB0739

**Fig. S6 Impaired microglial response to laser injury and decreased ramification index after PSB0739 treatment.** (a) Time-lapse images at the indicated time points showing microglial response to laser ablation in the indicated groups. Minutes post injury, mpi. Red lightning bolts indicate the ablation points. Scale bar, 20 µm. (b) Time-course of microglia response to laser ablation in different groups. 4 laser ablation in 3 mice in each group. Two-way ANOVA test for comparing response of Ctrl and PSB0739 group, ***P < 0.001. Error bar, SEM. (c) Statistics of microglial ramification index before and after injection of saline and PSB0739, respectively. Each line represents a cell and the bar shows the mean value. 15 microglia data from 3 mice in each group. Paired *t*-test, not significant (ns), ***P < 0.001.

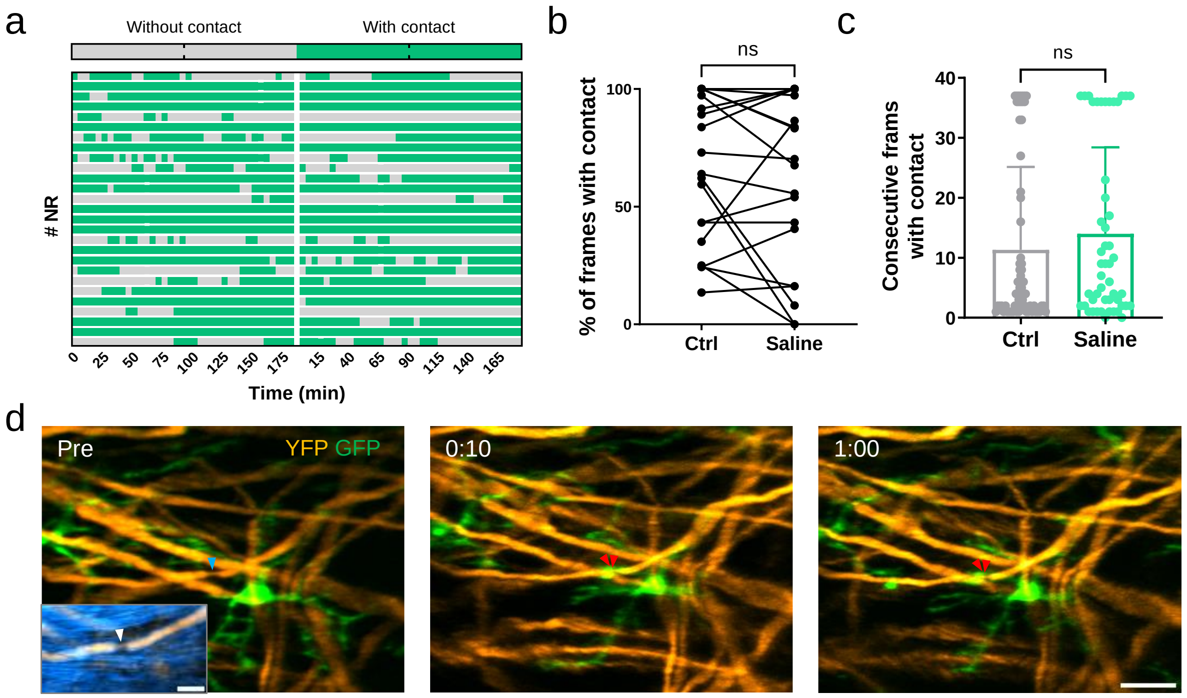


**Fig. S7 Intrathecal saline injection has no effect on physiological microglia-node interaction as well as microglial wrapping behaviour after axonal injury**. (a) Heating map showing microglia-nodes contacts during 3-h time-lapse imaging before and after saline treatment. The blank area indicates the time interval for saline injection. Images were taken every 5 minutes. A total of 27 nodes 3 mice were recorded. (b-c) Comparison of the percentage of frames with contact (b) and maximum contact lifetime (c) during 3-h time-lapse imaging before (Ctrl) and after saline treatment. Wilcoxon test for (b), Mann-Whitney test for (c). The statistics shown here are based on the data shown in (a). Error bar in (c) is standard deviation (SD). (d) Time-lapse images at indicated time points showing the interaction between axon (orange) and microglia (green) at NR following axonal injury with saline treatment. Images in (d) were representative of 5 axons. Blue arrowheads indicate the absence of contacts, double red arrowheads indicate the wrapping contacts. Insets, overlay of myelin SRS image (blue) and axon YFP image (orange). White arrowheads indicate the location of NR. Scale bars: 20 µm, 10 µm for the inset. Time is represented as hr: min.

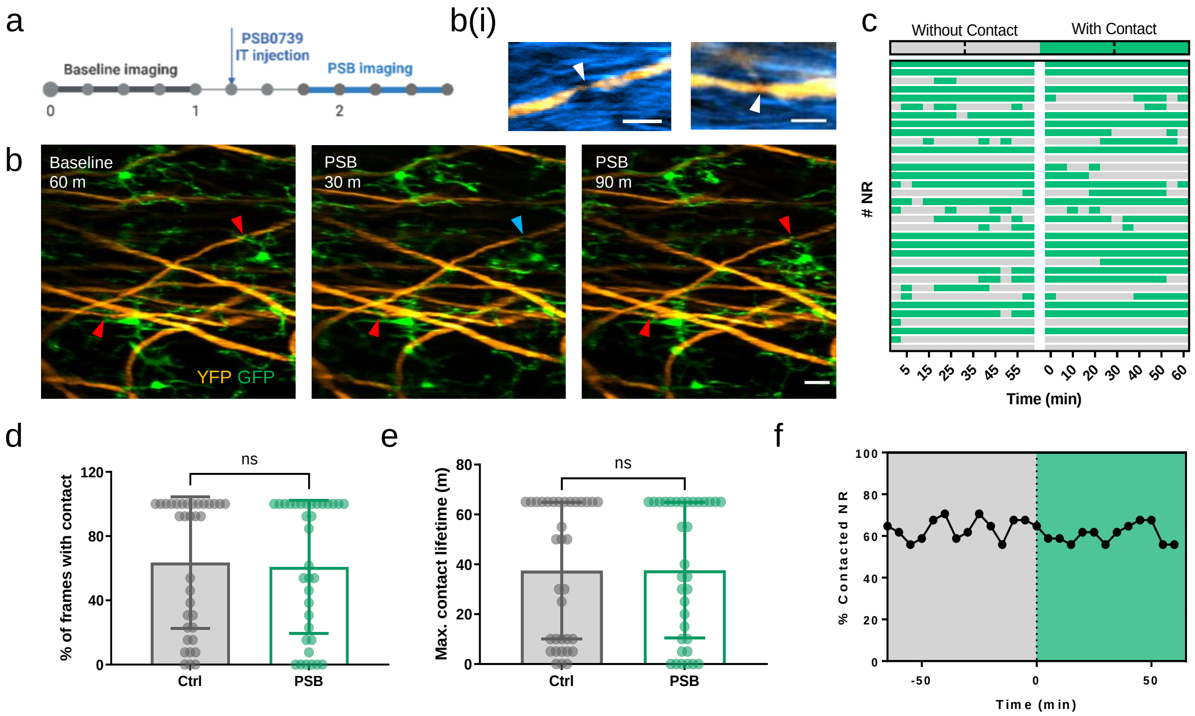


**Fig. S8 PSB0739 has no effect on microglia-node interaction under physiological condition.** (a) Illustration of the experimental design for the study of physiological microglia-node interactions with acute P2Y12 receptor blockade. IT, intrathecal. (b) Time-lapse images at indicated time points showing the physiological interaction between axon (orange) and microglia (green) at NR before and after PSB treatment. Red arrowheads indicate the occurrence of microglia-node contact, while blue arrowheads indicate the absence of contacts. Scale bar, 20 µm. b(i), merged SRS (blue) and YFP (orange) images of targeted axons showing the location of NR (white arrowheads) in (b). Scale bar, 10 µm. (c) Heatmap showing microglia-nodes contacts during 1-h time-lapse imaging before and after PSB treatment. The blank area indicates the time interval for PSB injection. Images were taken every 5 minutes. A total of 35 nodes from 3 mice were recorded. (d-e) Comparison of the percentage of frames with contact (d) and maximum contact lifetime (e) during 1-h time-lapse imaging before (Ctrl) and after PSB treatment. Error bars, SD. Mann-Whitney test. The statistics shown here are based on the data shown in (c). (f) The percentage of NR with microglial contact in the 35 recorded nodes showed little change after PSB treatment. The dashed line indicates the start point of PSB treatment.

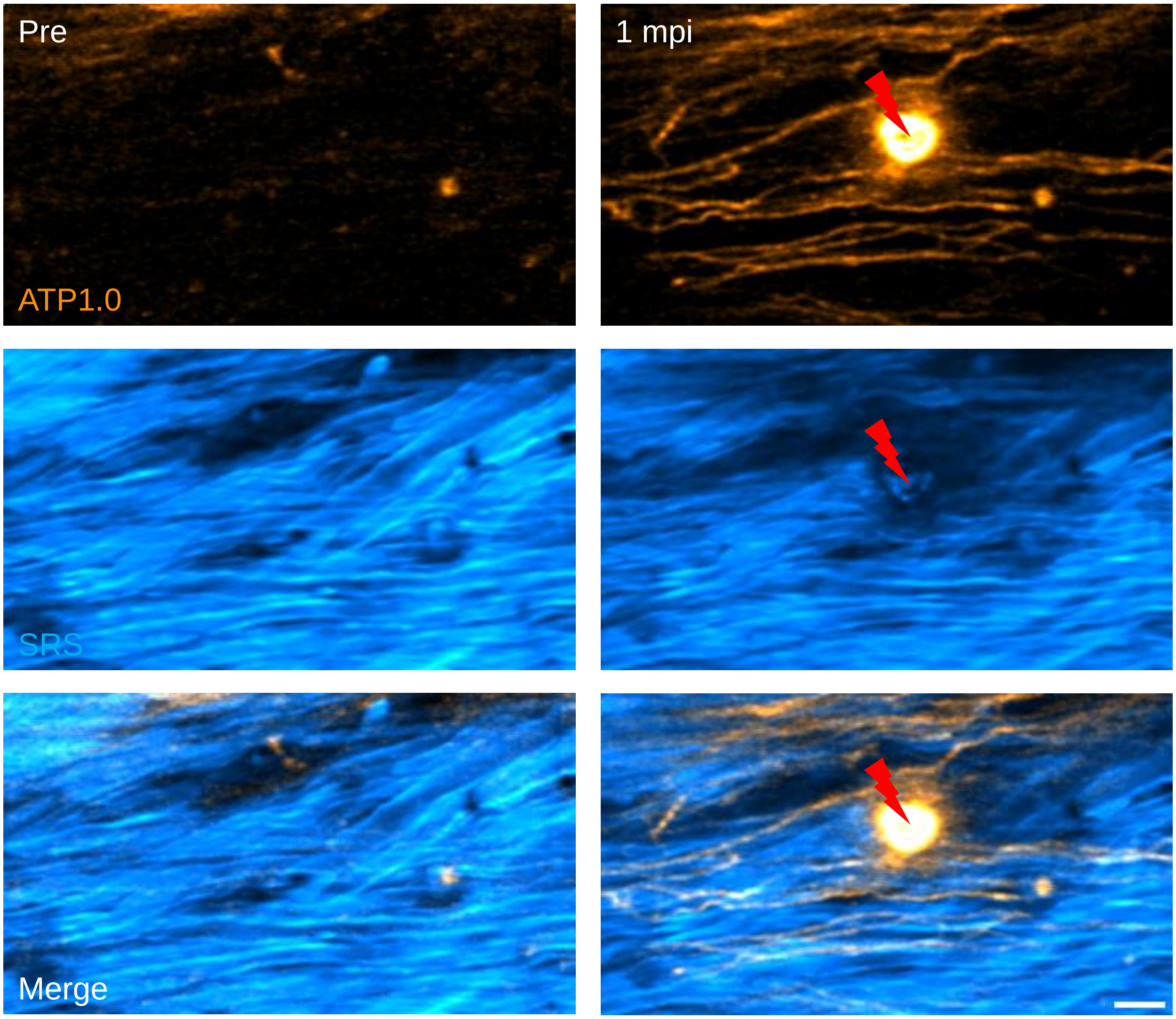


**Fig. S9. Only non-myelinated axons showed ATP fluorescent response to laser injury.** Time-lapse images at the indicated time points showing the ATP sensors' response to laser ablation. Prior to injury, the fluorescence of ATP sensors was weak. After a local laser injury, the ATP sensors fired immediately. Red lightning bolts indicate the ablation points. Orange, ATP1.0; blue, SRS signal of myelin sheath. Scale bar, 10 μm.

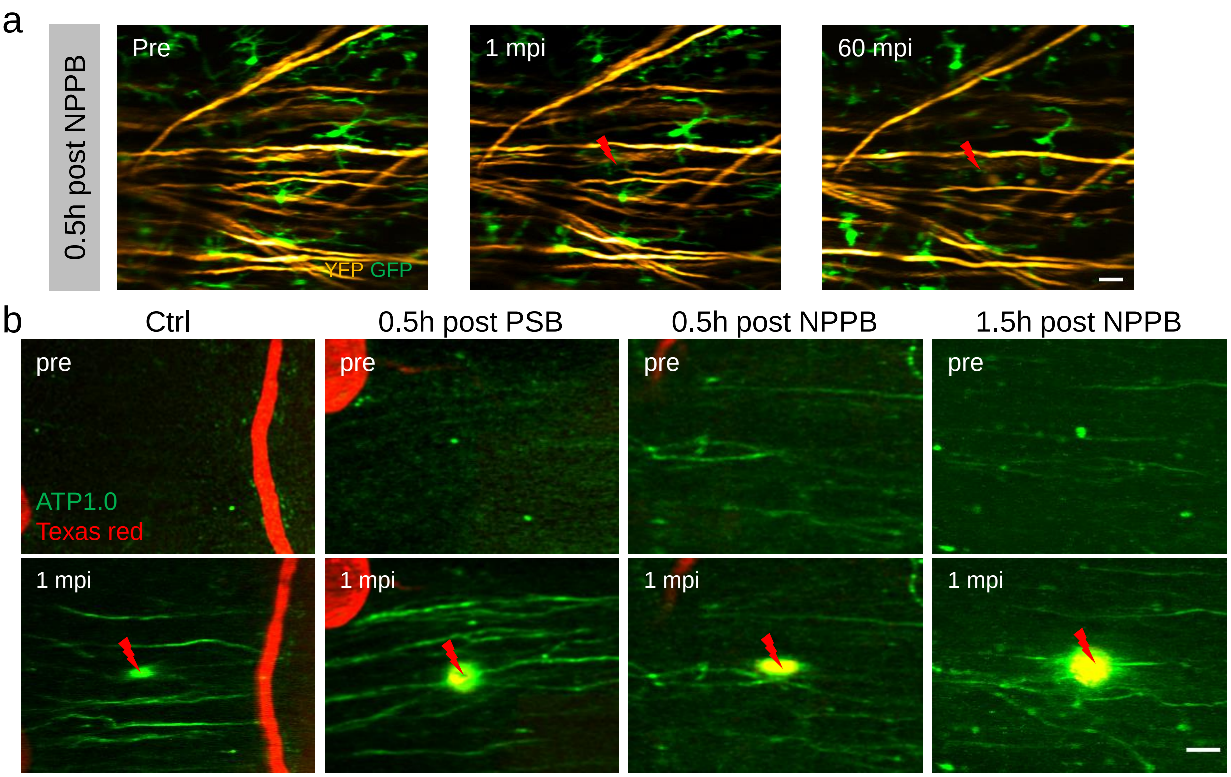


**Fig. S10 Impaired microglial directed motility and ATP release in response to laser injury after NPPB treatment.** (a) Time-lapse images show microglial response to laser ablation at 0.5 hr post NPPB injection. Orange, YFP-labeled axon, green, GFP-labeled microglia. (b) Representative images of the ATP sensor response to the laser injury in the indicated groups. NPPB treatment impaired the response of ATP sensor significantly. Red lightning bolts indicate the ablation points. Green, ATP1.0 labeled axons, red, Texas red labeled blood vessels. Scale bars, 20 μm in (a-b).

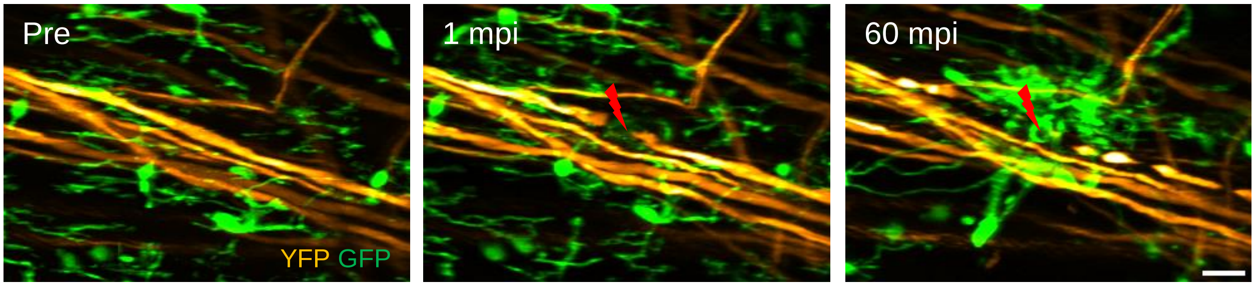


**Fig. S11 TTX has no effect on the microglial response to laser injury.** Time-lapse images at the indicated time points show the response of microglia (green) to laser injury of an axon (orange) in the TTX treated group. Red lightning bolts indicate the ablation points. Scale bar, 20 μm.

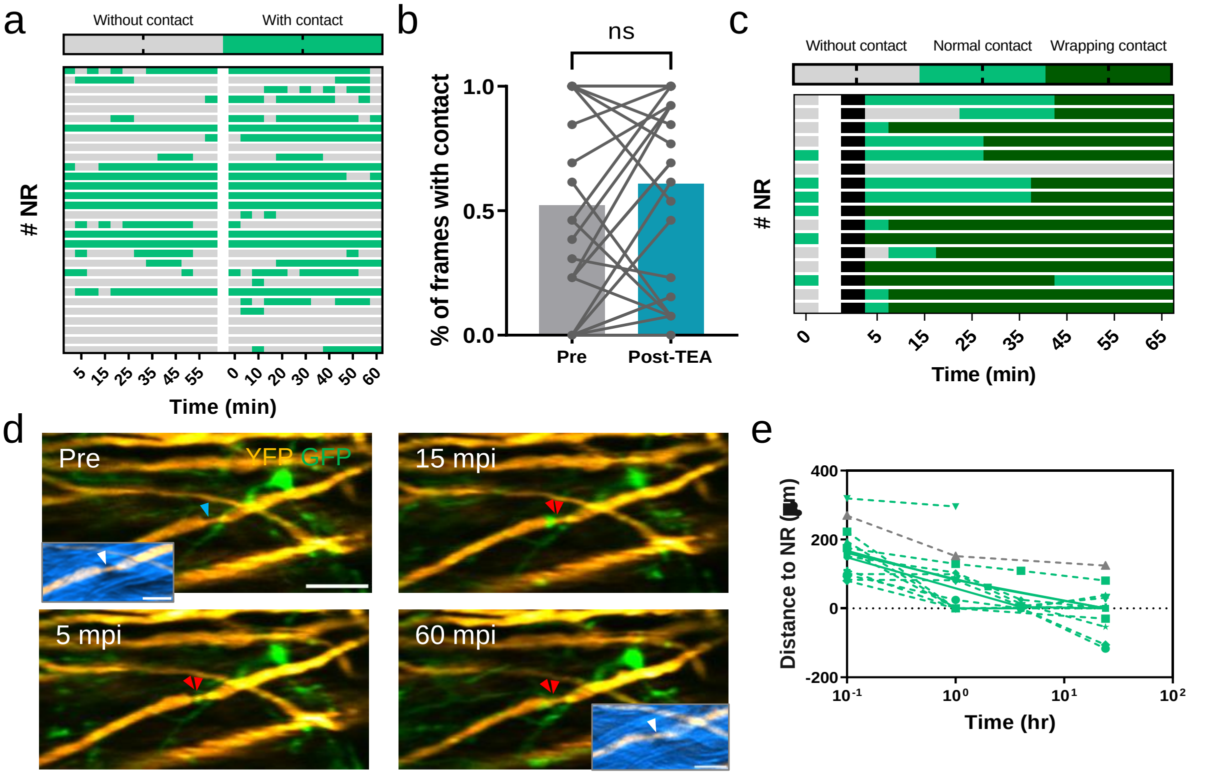


**Fig. S12 TEA has no effect on the microglia-node interaction under both physiological and pathological conditions.** (a) Heatmap showing microglia-nodes contacts during 1-h time-lapse imaging before and after TEA treatment. The blank area indicates the time interval for TEA injection. Images were taken every five minutes. A total of 40 nodes from 4 mice were recorded. (b) Comparison of the percentage of frames with contact during 1-h time-lapse imaging before (Ctrl) and after TEA treatment. Mann-Whitney test. (c) Heatmap showing microglia-node contacts from 5 mins before to 1 hr of axon after laser axotomy in the TEA treated group. The blank area indicates the interruption of time-lapse imaging when laser injury was performed. Images were taken every 5 minutes. A total of 16 axons from 6 mice were recorded. (d)

Time-lapse images showing the microglia-node interaction at the indicated time points before and after injury. Blue arrowheads indicate nodes without microglial contact and red arrowheads indicate nodes contacted by microglia (green). Double red arrowheads indicate wrapping contact at NR. Insets, overlay of myelin SRS image (blue) and axon YFP image (orange). White arrowheads indicate the location of NR. (e) The distance between the proximal ends of injured axons and the proximal NR over the first 24 hr after injury in the TEA-treated group. The grey color indicates the data of axons without wrapping contact. Curves across the dashed line indicate the axonal degeneration events with nodal breach. Scale bars, 20 μm in (d), 10 μm for all the insets.

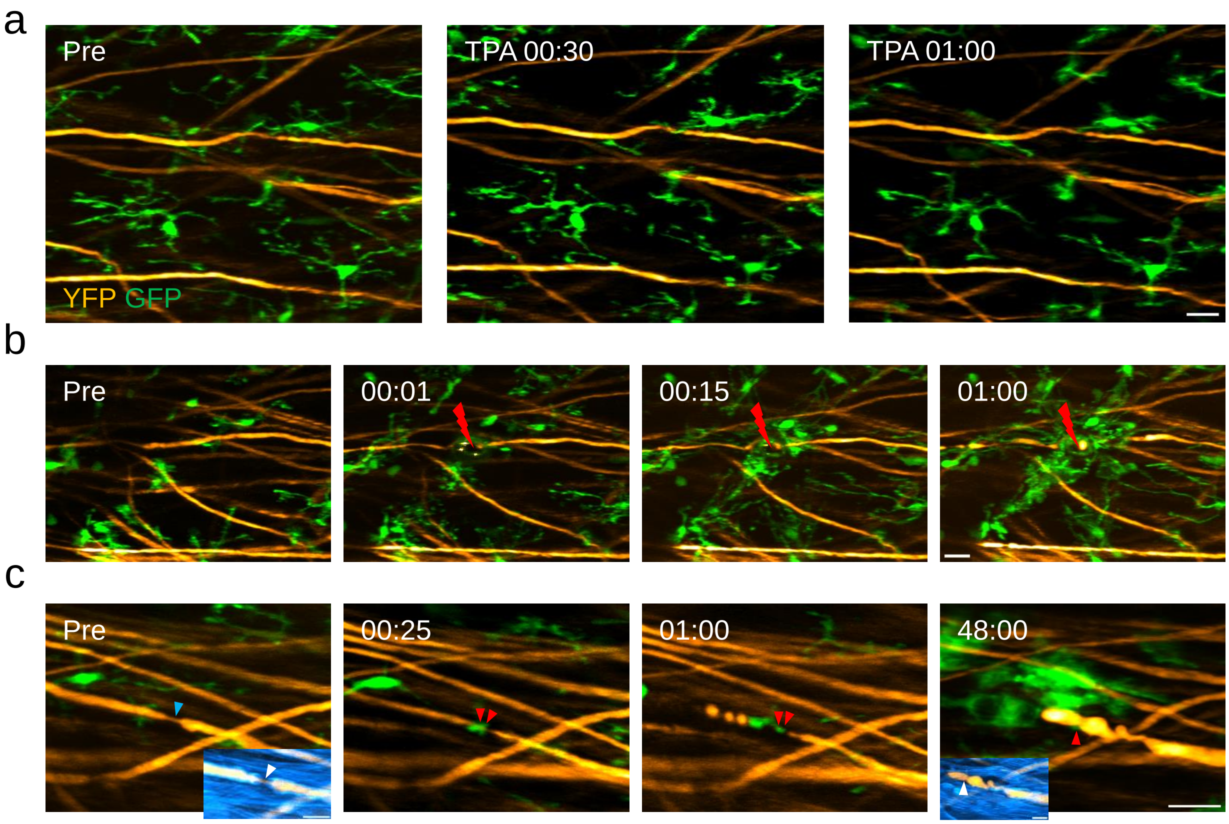


**Fig. S13 TPA has no effect on the microglial wrapping contact with NR after axonal injury.** (a) Time-lapse images at the indicated time points showing the effect of TPA on microglia ramification. Orange, YFP-labelled axons; green, GFP-labelled microglia. (b) Time-lapse images at the indicated time points showing the response of microglia after laser ablation in the presence of TPA treatment. Red lightning bolts indicate the ablation points. (c) Time-lapse images at indicated time points showing the interaction between axon (orange) and microglia (green) at NR after injury with TPA treatment. Blue arrowheads indicate the absence of contacts, double red arrowheads indicate the wrapping contacts. Insets, overlay of myelin SRS image (blue) and axon YFP image (orange). White arrowheads indicate the location of NR. Scale bars: 20 μm in (a-c), 10 μm for all the insets in (c). Time is represented as hr: min.
